# A novel CHD4-CTCF regulatory axis drives key developmental programs and transitions in male germ cells

**DOI:** 10.1101/2025.09.26.678824

**Authors:** Rodrigo O. de Castro, Agustin Carbajal, Katarzyna P. Nowak, Ayelén González Montoro, Monika Kawecka, Luciana Previato, Roberto J. Pezza

**Affiliations:** Cell Cycle and Cancer Biology Research Program, Oklahoma Medical Research Foundation, Oklahoma City, Oklahoma, United States of America; Doctoral School of Molecular Biology and Biological Chemistry, Warsaw, Poland; Department of Cell Biology. University of Oklahoma Health Sciences Center, Oklahoma City, Oklahoma, United States of America

**Author notes:** These authors contributed equally to this work (Rodrigo O. de Castro, Agustin Carbajal). Corresponding author: Roberto J. Pezza. Suite B305. 825 NE 13^th^ street, Oklahoma City, Oklahoma, 73104.

**Keywords:** mouse gametogenesis, spermatogonia, meiosis, CTCF, CHD4, and chromatin remodeling

## Abstract

Sustained gamete production and testis development relies in developmental programs and transitions such as spermatogonia differentiation, meiosis, and spermiogenesis. Chromatin remodeling complexes and CTCF are key regulators of these processes by controlling specific patterns of gene expression across gamete development. Although the CHD4 chromatin remodeler and CTCF have been implicated in spermatogonia development, a mechanism explaining their cooperation is missing. We also do not know CHD4 and CTCF role in later stages of spermatogenesis. Our studies reveal that germ cell-specific deletion of CHD4 results in spermatogonia development defects and affects meiotic-specific functions such as homologous chromosome synapsis. Combined analysis of genome-wide chromatin binding and single cell gene transcription showed that CHD4 controls specific gene expression programs important for spermatogonia differentiation, transition into meiosis, and meiotic specific processes. Mechanistically, this can be explained by a pervasive role for CHD4 in controlling CTCF chromatin binding and transcriptional function. The discovery of a functional interplay between CHD4 and CTCF reveals a novel axis of gene expression control, integrating chromatin structure and transcriptional outcomes. This interaction is essential in orchestrating early testis development and meiotic processes, ensuring that the correct number of chromosomes is faithfully transmitted to the next generation.

## Introduction

Defects in gametogenesis leading to the production of aneuploid gametes are a major cause of birth defects and infertility. Aneuploidy also poses a significant challenge for couples undergoing assisted reproductive technologies, where over 30% of preimplantation embryos are aneuploid. Additionally, understanding the mechanisms underlying germ cell development is crucial for improving outcomes in common gonadal developmental disorders.

Male germ cell development follows a sequential differentiation pathway that begins with development of spermatogonial stem cells followed by their commitment to meiosis^1^ and spermiogenesis, ensuring the proper development of functional sperm^2^. In meiosis, coupled to homologous recombination-dependent repair of DNA double strand breaks, the homologous chromosomes undergo chromosome pairing and synapsis. This is mediated by the synaptonemal complex, which starts to assemble at the prophase stage of leptotene and is fully formed by pachytene. Cells remain in pachytene until re-combination and synapsis checkpoints are satisfied, after which the synaptonemal complex disassembles as the meiotic cell progresses through diplotene.

Chromatin undergoes extensive remodeling during gametogenesis, leading to altered gene expression and chromosome organization^3,4^. This remodeling is proposed to control obligatory developmental transitions such as the conversion from undifferentiated to differentiated spermatogonia, spermatogonia commitment to meiosis, and meiotic exit.

CHD4 (chromodomain helicase DNA-binding protein 4) is an ATP-dependent chromatin remodeler member of the SNF2 superfamily of ATPases that regulates genome function by repositioning nucleosomes and altering local chromatin structure^5–7^. Through its remodeling activities, CHD4 modulates the accessibility of transcription factors and DNA repair machinery to specific regions of the chromatin, thereby influencing transcriptional activity, DNA damage repair, and the preservation of genome stability. Recent studies have explored the roles of CHD4 during premeiotic stages of spermatogenesis^8–10^. Germ cell-specific deletion of *Chd4* leads to arrested early gamete development due to the failed survival of neonate undifferentiated spermatogonial stem cells^9^. Gene expression analyses following *Chd4* knockdown have shown reduced expression of genes associated with spermatogonial self-renewal and increased expression of progenitor cell signature genes^8^. CHD4 also establishes chromatin states required for the formation and maintenance of the ovarian reserve and closed chromatin states at the regulatory elements of pro-apoptotic genes in male germ cells, allowing male germ cell survival^11^.

CHD4 is a fundamental component of the NURD (NUcleosome Remodeling and Deacetylase), a pivotal chromatin-modifying complex that regulates gene expression through chromatin remodeling and histone deacetylation. NURD is crucial for various developmental and somatic cell processes, including stem cell differentiation, cell identity maintenance, and DNA damage response^5–7^. CHD4 also interacts with ADNP and HP1 to from the ChAHP complex, which has been implicated in critical cell functions such as lineage gene expression in ESCs^12^. Recently, it has been shown that ADNP regulates local chromatin architecture by competing for binding with CTCF, a master genome architecture protein^13^.

In this study, we generated a novel testis specific knockout model allowing studies of CHD4 deletion in later spermatogonia and meiotic spermatocytes. We demonstrate that germ cell–specific deletion of CHD4 leads to deficient spermatogonia development and disrupted meiotic progression, with mutant primary spermatocytes arrested at a zygotene-like stage and undergoing increased apoptosis. CHD4-deficient spermatocytes exhibit moderate defects in DNA repair, incomplete synaptonemal complex assembly, and defective homologous chromosome synapsis. Genome-wide chromatin binding and single cell transcriptional analyses reveal mechanistic insights into CHD4 function. Specifically, CHD4 regulates CTCF chromatin binding and transcription programs in spermatogonia and meiotic spermatocytes. Our results define a CHD4–CTCF regulatory axis essential for normal testis development and the production of functional gametes.

## Results

### CHD4 is required for spermatogonia development and completion of meiosis

Previous studies using Ddx4-Cre mice have demonstrated that CHD4 is essential in premeiotic phases of gamete development^9,10^, making it unfeasible to study possible functions of CHD4 later during meiosis using this model. Thus, we generated *Chd4*^−/−^ mice using Ngn3-Cre, which is activated in differentiating spermatogonia^14^ and thus it is suited for gene deletion at meiotic prophase I and later (Fig. 1A). Successful recombination of loxP sites in mouse testis was confirmed by genotyping PCR (Fig. 1B). CHD4 protein levels were strongly reduced in *Chd4^−/−^*testis evaluated by western blot of 2 month old mice whole testis extract (Fig. 1C) or immunostaining of paraffin-embedded testis sections (Fig. 1D), indicating that this is a null allele.

**Figure 1.**
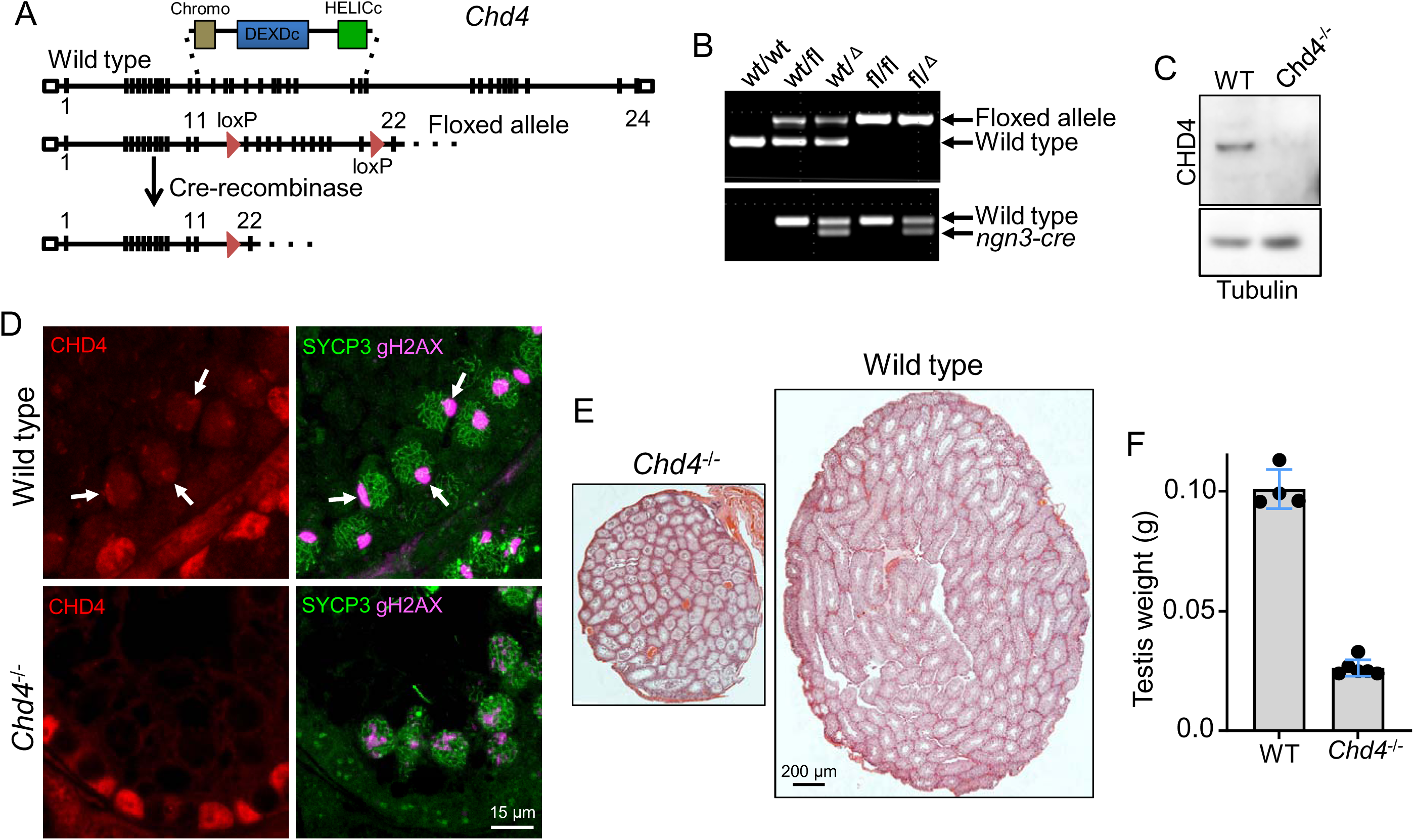
Impaired gamete development in *Chd4*^−/−^ testis. **(A)** *Chd4* gene targeting design. **(B)** Genotyping PCR showing *Chd4* wild type and floxed alleles and Ngn3 positive mice. **(C)** Western blot showing CHD4 depletion in whole testis extract of 2 months old *Chd4^−/−^* testis. **(D)** Expression of CHD4 in wild type and *Chd4^−/−^* testis assessed by immunofluorescence. Arrow indicates the presence of CHD4 at the sex body in pachynema (SYCP3/yH2AX staining). Arrowhead indicates the absence of CHD4 in zygotene-like cells of *Chd4^−/−^* testis. Similar results were observed in 3 different mice. **(E)** H&E-stained histological sections of wild type and *Chd4*^−/−^ testes. **(F)** Quantitation of wild type and *Chd4*^−/−^ testis size. Compare mutant 0.026 ± 0.003, n = 6 mice (mean ± SD) with wild type 0.100 ± 0.008, n = 4 mice; P < 0.0001 (two-tailed Student t test).

*Chd4^−/−^* adult mice (2 months old) appeared normal in all aspects except in the reproductive tissues. Testes were significantly smaller in *Chd4^−/−^* males (mean: 0.026g ± SD: 0.003, number of quantified mice nLJ=LJ4 (8 testes), P ≤ 0.0001, t test) compared to wild type (0.10 g ± 0.008, nLJ=LJ6 mice (12 testes)) littermates (Fig. 1 E and F), indicating severe developmental defects in the testis.

To assess male meiosis I defects in detail, we analyzed H&E-stained testis sections from 45 days wild type and *Chd4^−/−^* mice (Fig. 2A). This analysis revealed that gametogenesis progresses trough initial stages of prophase I, but cells are arrested in a zygotene-like stage (Fig. 2A), consistent with an effect in meiotic stages of development. The *Chd4^−/−^* mutant testes showed signs of substantial apoptosis in seminiferous tubules Fig. S1A). A meiotic phenotype was confirmed by scoring individual stages of asynchronous populations of wild type and *Chd4^−/−^* spermatocytes from 2 month old mice (Fig. S1B). Here, we observed a significant relative increase in zygotene cells in *Chd4^−/−^* versus wild type mice and a decrease in pachytene and diplotene spermatocytes when *Chd4^−/−^* mice is compared to wild type (Fig. S1B).

**Figure 2.**
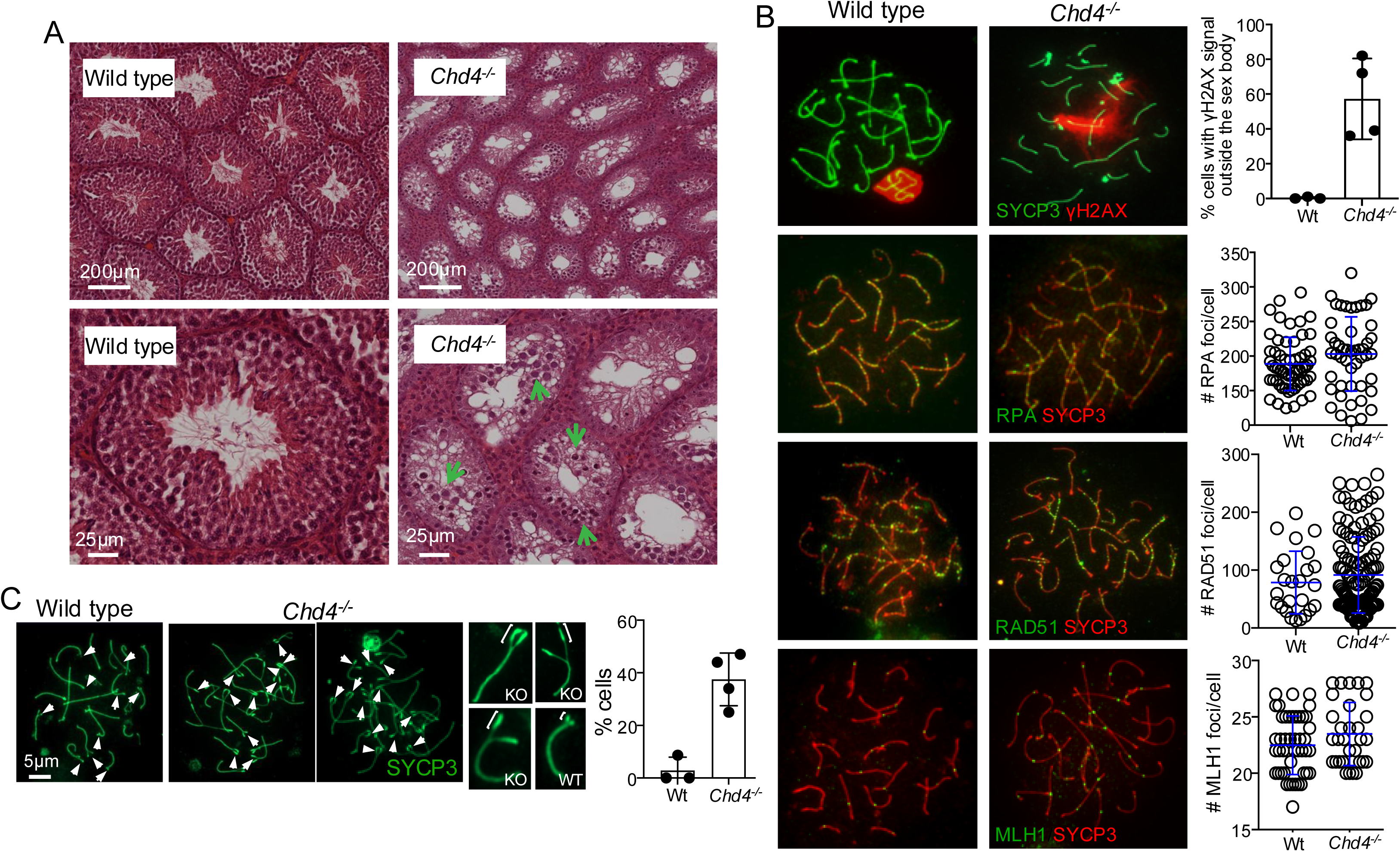
*Chd4*^−/−^ mice show profound defects in gametogenesis. **(A)** H&E-stained histological sections of wild type and *Chd4*^−/−^ testes. Green arrows indicate the most advanced cells in *Chd4^−/−^* mouse. **(B)** Representative images of wild type and *Chd4^−/−^* spermatocytes immunostained with SYCP3 and γH2AX, RPA, RAD51, or MLH1. Quantification of percentage of cells with γH2AX signal outside the sex body in the most advanced spermatocytes is also shown (wild type 2 ± 1, *n* = 62 cells, 3 mice; *Chd4^−/−^* 58 ± 11, *n* = 48, 3 mice, P = 0.0001). Quantification of the number of foci per cell for RPA (wild type 188.6 ± 38.6, *n* = 58 cells, 3 mice; *Chd4^−/−^* 203.1 ± 53.7, *n* = 48, 3 mice, P = 0.11), RAD51 (wild type 78.6 ± 54.08, *n* =25 cells, 3 mice; *Chd4^−/−^* 91.65 ± 65.81, *n* = 127, 3 mice, P = 0.35), and MLH1 (wild type 22.5 ± 2.6, *n* = 46 cells, 3 mice; *Chd4^−/−^* 23.5 ± 2.8, *n* = 31, 3 mice, P = 0.11) is also shown. **(C)** CHD4 deletion results in abnormal homologous chromosome synapsis. Representative images of wild type and *Chd4^−/−^* spermatocytes immunostained with SYCP3. White arrows mark unsynapsed portions of chromosomes. The insets on the right show higher magnification of SYCP3 images.

We also analyzed H&E-stained histological sections of ovaries from 2-month-old wild type and *Chd4^−/−^* female mice. We noted no significant difference regarding ovary size, changes in stromal cells, or reduced number of follicles in the mutant with respect to wild type (Fig. S1C).

In sum, our results indicate that CHD4 has a dual function in spermatogenesis. independent of its role in spermatogonia^8–10^, CHD4 is required for meiotic progression of mice male germ cells.

### Analysis of cytological markers of DSB repair in *Chd4^−/−^* and wild type spermatocyte

CHD4 is known to play a role in the DNA damage response in somatic cells^16,17^. For example, *Chd4* knockdown results in increased γH2AX, a marker of double-strand breaks, and CHD4 accumulates at sites of DNA damage^15^. To determine whether CHD4 is required to repair the programmed DSBs that occur in germ cells during meiosis, we immunostained spermatocyte chromosome spreads for γH2AX, a marker of DNA damage. In agreement with a mid-meiotic arrest, we observed γH2AX immunostaining outside the sex body in a fraction of *Chd4^−/−^* spermatocytes compared to wild type (57.2 ± 23.2%, n = 4 mice versus 0.33 ± 0.57%, n = 3 mice; mean ± SD; P = 0.009, two tailed t test) (Fig. 2B). The abnormal distribution of γH2AX signal we observed suggests a moderately deficient repair of DSBs.

Next, we measured foci number of later markers of DNA repair, RPA, RAD51, and MLH1. RPA binds to partial single strand DNA sites product of DNA end resection, then RAD51 interacts with resected DSBs displacing RPA from these sites and catalyzes strand invasion^18^. MLH1 is involved in resolution of DNA branched structures and is a marker of crossover sites^19^. Comparative analysis between wild type and *Chd4^−/−^* spermatocytes shows no significant differences in number of foci per cell for RPA (188.6 ± 38.5%, n = 98 cells from 3 mice versus 203.1 ± 53.7%, n = 85 cells from 3 mice; mean ± SD; P = 0.03, two tailed t test), RAD51 (78.6 ± 54.1%, n = 25 cells from 3 mice versus 91.7 ± 65.8%, n = 126 cells from 3 mice; mean ± SD; P = 0.35, two tailed t test), or MLH1 (22.5 ± 2.6%, n = 46 cells from 3 mice versus 23.5 ± 2.8%, n = 31 cells from 3 mice; mean ± SD; P = 0.11, two tailed t test) (Fig. 2B).

We conclude that *Chd4^−/−^*spermatocytes may have a mild DNA repair deficiency, suggested by abnormal γH2AX kinetics.

### *Chd4^−/−^* spermatocytes exhibit defects in homologous chromosome synapsis

During meiotic prophase, the association of the homologous chromosome pairs is stabilized by the synaptonemal complex. Homologous synapsis can be monitored by following the immunosignal of proteins of the lateral element (e.g., SYCP3) and the central region (e.g., SYCP1) on chromosome spread preparations. Homologous pairing and initiation of synapsis appeared normal in *Chd4^−/−^* spermatocytes, as judged by staining patterns for SYCP3 and SYCP1 in leptotene and zygotene cells, but proper pachynema with fully synapsed autosomal bivalents was rarely observed (Fig. 2C). Instead, a good portion of the most advanced cells showed a late zygotene-like morphology (full-length axes have been developed and synapsis is near complete) but with characteristics of pachytene cells such as acquisition of the characteristic knob-like accumulation of SYCP3 at telomeres (Fig. 2C, and Fig S1D). A detailed analysis of *Chd4^−/−^* spermatocytes revealed abnormal formation of the synaptonemal complex at pericentromeric areas with notable over deposition of SYCP3 (wild type 2.90 ± 5.02 % of cells, n = 3 mice versus *Chd4^−/−^* 37.50 ± 10.01 %, n = 4 mice; mean ± SD; P = 0.0017, two tailed t test). These abnormal pericentromeric areas are devoid of both SYCP1, marking transverse filaments, and SYCE1, from the central element of the synaptonemal complex (Fig. S1D and E).

We conclude that *Chd4^−/−^*spermatocytes exhibit incomplete synapsis associated with defects in the synaptonemal complex assembly and/or maturation, a characteristic that likely triggers cell arrest and apoptosis.

### CHD4 primarily acts as a transcriptional repressor during spermatogonia development and early meiosis

As a chromatin remodeling factor, CHD4 likely influences gene transcription during spermatogenesis. To directly test CHD4-mediated cell type specific transcriptional changes, we performed single cell RNA-seq using 60 days mice old testis. Two *Chd4^−/−^* and two wild type littermate mice were processed in parallel and analyzed using 10 X Genomics 3’ single cell platform. After filtering out empty droplets, doublets, and low quality cells, we obtained a total of 12,166 cells evenly distributed between wild type (6,671) and *Chd4^−/−^* (5,495) cells. General quality control metrics of the final dataset can be seen in Figure S2. Unsupervised clustering of cells resulted in 21 different cell clusters (Fig. S3A).

To determine cell types present in each cluster we used a combination of two complementary strategies: (1) label transfer using the scRNA-seq dataset from Chen et al., 2018^20^ as reference (Fig. S3B and C), and (2) analysis of conventional marker gene expression (Fig. S3D) The final labels assigned to each cell cluster can be seen in Fig. S3A, together with a plot showing the distribution of cells per cluster and genotype (Fig. S3E).

Our dataset contains 13 germ cell (GC) clusters (GC_A to GC_M) (Fig. S3A), representing the whole spectrum of germ cells, from spermatogonia, characterized by the expression of gene markers such as *Zbtb16*, *Dmrt1*, *Stra8*, *Gfra1*, and *Sall4*, to elongating spermatids, characterized by the expression of genes such as *Tppp2*, *Prm2*, and *Prm1* (Fig. S3D, Table S1). Our data also includes somatic cells with a large number of Sertoli cells (expressing *Sox9*), Leydig cells (expressing *Insl3*), and a minor number of macrophages (*Lyz2*), endothelial cells (*Flt1*) and fibroblast/myoid cells (*Col1a2*) (Fig. S3A, E). Compared to wild type, germ cell development is arrested in *Chd4^−/−^* mice. This is reflected in a reduced number of cells from primary spermatocytes to elongated spermatids and an increased number of spermatogonia and somatic cells (Fig. S3A, E).

Given that *Chd4^−/−^* germ cells rarely differentiate past pachynema, we focused our analysis on the early germ cell stages encompassing spermatogonia through pachytene. To gain resolution for these stages, we re-processed this subset of germ cells (Fig. 3A-C), corresponding to clusters GC_A to GC_C (spermatogonia to pachytene, Fig. S3A). This resulted in a new dataset of 2,135 cells (854 wild type and 1,281 *Chd4^−/−^* cells) grouped into five clusters, which we designated as GC_1 to GC_5 (Fig. 3A and B).

**Figure 3.**
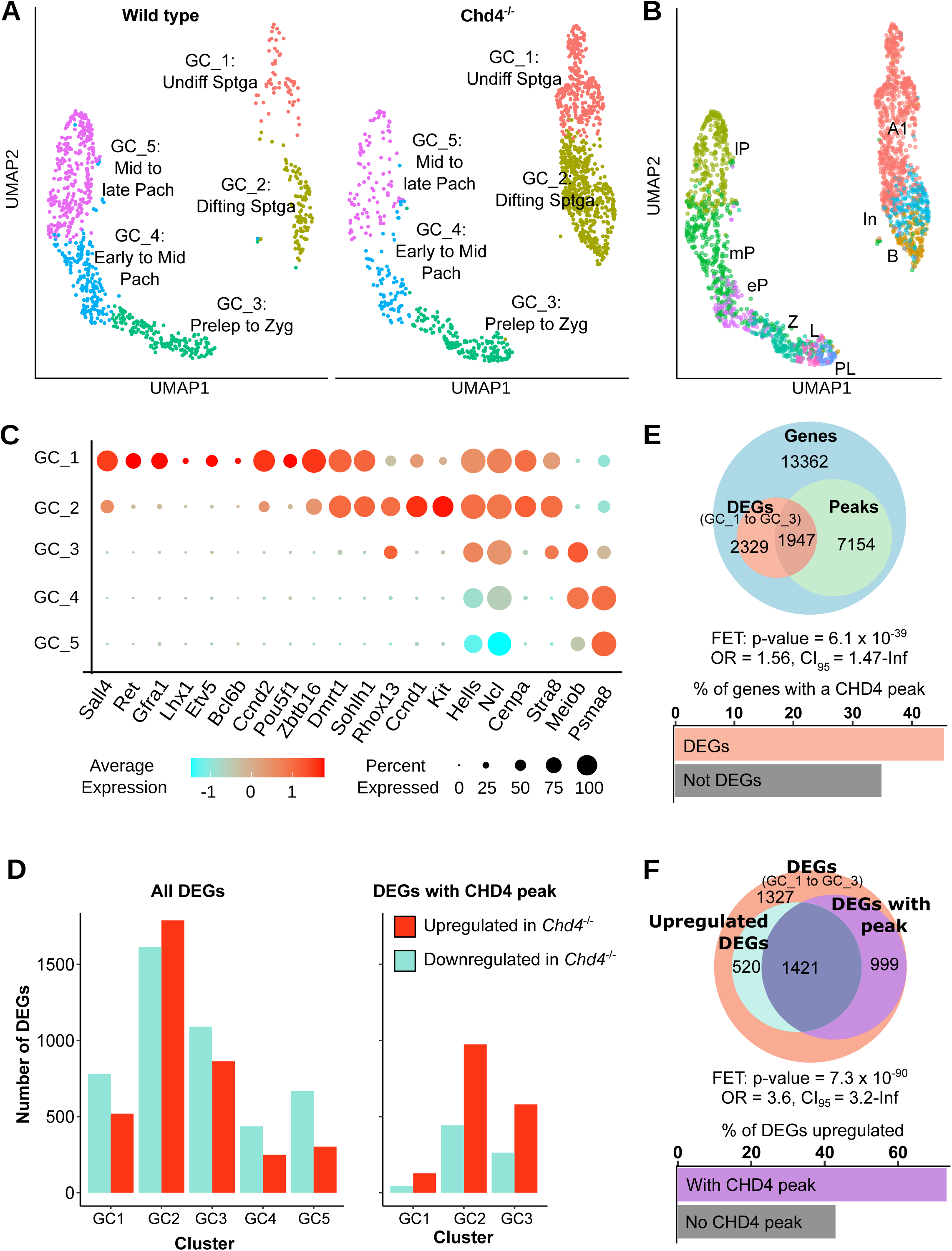
Analysis of scRNA-seq early germ cell subset. CHD4 has a predominately repressive effect. **(A)** Clustering of early germ cells scRNA-seq experiment. Germ cells from spermatogonia to pachytene from the scRNAseq experiment shown in Supplemental Figure 4A were reprocessed as an independent group. The graphs show these cells in UMAP space colored by cluster and split by genotype. **(B)** Cell label transfer using the dataset from Chen et al, 2018. To estimate the identities of cells in our dataset we performed label transfer using the dataset from Chen et al, 2018. This dataset is a scRNA-seq performed on synchronized and purified germ cells from spermatogonia A to rounded spermatids. Cells’ labels are as follows: spermatogonia A1 (’A1’), Intermediate spermatogonia (’In’), spermatogonia B (’B’), preleptotene (’PL’), leptotene (’L’), zygotene (’Z’), early pachytene (’eP’), middle pachytene (’mP’) and late pachytene (’lP’). **(C)** Expression of germ cell markers per cluster. Dot plot depicting the expression of selected marker genes across clusters. The color indicates scaled average expression (z-score) and the dot size represents the percentage of cells expressing the gene in the cluster. **(D)** CHD4 has a predominately repressive effect on gene expression. Left: Bar plot depicting the number of DEGs per cluster that increase (red) or decrease (blue) their expression in *Chd4^−/−^*with respect to wild type mice. Right: same as left but only counting DEGs annotated with a CHD4 peak. Cluster to peak-set matching criterion is the same as for (D). **(E)** Genes that change expression in the *Chd4^−/−^* mice from spermatogonia to zygotene are associated to genes bound by CHD4. Top: Euler diagram showing the overlap between genes that are differentially expressed in clusters GC_1, GC_2 or GC_3 (’DEGs (GC1 to GC3)’) and genes that were annotated with a CHD4 consensus peak in spermatogonia or preleptotene peak sets (’Genes associated with a CHD4 peak’), among all genes detected in the scRNA-seq experiment (’Genes’). DEGs from GC_1 were matched to genes annotated with a spermatogonia peak, DEGs from GC_2 were matched to genes annotated with a spermatogonia or preleptotene peak, and DEGs from GC_3 were matched to genes annotated with a preleptotene peak. Middle: p-value, odds ratio (OR) and confidence interval (95%) calculated with two-sided Fisher’s exact test (FET), for categories ‘gene is differentially expressed in GC_1, GC_2 or GC_3’ vs ‘gene is annotated with CHD4 peak’ (from euler’s plot above). Bottom: bar plot depicting percentage of genes that have a CHD4 peak associated to them, for genes that differentially expressed in GC_1, GC_2 or GC_3 (’DEGs’) or not (’Not DEGs’). **(F)** DEGs bound by CHD4 are associated to upregulated DEGs. For clusters GC_1, GC_2 and GC_3, Top: Euler diagram showing total number of genes that are differentially expressed in any of the clusters (’DEGs (GC1 to GC3)’), number of DEGs that increase their expression in *Chd4^−/−^*(’Upregulated DEGs’) and DEGs that have a CHD4 peak annotated to them. Criteria to associate a cluster to a peak set is the same as for (D). Middle: p-value, odds ratio and confidence interval (95%) from Fisher’s exact test, on categories ‘DEG has a CHD4 peak annotated to it’ and ‘DEG has a higher expression in *Chd4^−/−^*’ for all genes differentially expressed in above mentioned clusters. Nine genes that are upregulated in one cluster and downregulated on another were excluded from the analysis. Bottom: percentage of DEGs from above mentioned clusters that increase their expression in *Chd4^−/−^*, for DEGs with or without a CHD4 peak annotated to them. Same genes as before were excluded.

Cluster GC_1 is mostly composed of undifferentiated spermatogonia, expressing genes such as *Sall4*, Ret, *Gfra1*, *Lhx1*, *Bclb6*, and *Pou5f1* while having little expression of differentiating spermatogonia genes such as *Kit*. While label-transfer using the dataset from Chen data labels most of GC_1 cells as A1 spermatogonia, this is not an accurate assignation because A1 is the most undifferentiated label present in the reference dataset (Fig. 3B and S3D). GC_2 cells consist of differentiating spermatogonia. They have high expression of *Kit*, low expression of typical undifferentiated spermatogonia genes, and low expression of meiotic specific genes such as *MeioB.* Consistently, label-transfer assigns them the spermatogonia A1, intermediate, and B labels (Fig. 3C and Fig. S3F). Cells in GC_3 show high expression of typical leptotene/zygotene genes such as *MeioB* and *Dmc1*, and low expression of spermatogonia genes (*Kit*) as well as pachytene genes such as *Piwil1* and *Psma8* (Fig. 3C). The label-transfer approach identifies most cells in this group as in zygonema, while also containing a minor number of pre-leptotene, leptotene and early pachytene cells (Fig. S3F). Therefore, cells in the GC_3 cluster are labeled as pre-leptotene to zygotene (Fig. 3A). Finally, GC_4 and GC_5 cells, correspond to pachynema by label transfer, and accordingly, express typical pachytene cell genes.

To gain insight into cell type-specific gene regulation by CHD4 we analyzed differential gene expression for each cell cluster in the *Chd4^−/−^* mouse. We detected a total of 4,552 differentially expressed genes across all five clusters, which comprise about 18% of all detected genes. These genes are evenly distributed between upregulated and downregulated genes (Fig. 3D, left panel, Table S2).

To assess which genes are directly regulated by CHD4 we used ChIP-seq and CUT&RUN data. To assess CHD4 binding regions in spermatogonia cells we datamined four experiments performed using 7 dpp mice (^9,10^, see Table S3 for details). To assess CHD4 binding regions in preleptotene/leptotene cells, we performed two CUT&RUN experiments using WIN18,446/retinoic acid-synchronized testes, at 7 days post-retinoic acid injection, which results in enrichment of a spermatocyte population at the preleptotene/leptotene stages. Consensus regions obtained from replicate samples were designated as ‘spermatogonia’ or ‘preleptotene/leptotene’ peak sets, respectively, and peaks were annotated to the gene with the closest transcription start site (Fig. S4). Annotated spermatogonia peaks were matched to differentially expressed genes (DEGs) from clusters GC_1 or GC_2, while preleptotene/leptotene peaks were matched to DEGs from clusters GC_2 and GC_3. We observed that 45% of DEGs are associated with a CHD4 peak, which is a statistically significant association (Fig. 3E) (Fisher’s exact test (FET), p-value < 6×10^-39^, odds ratio (OR) = 1.56), suggesting that CHD4 regulates transcription of genes by direct action. Importantly, the majority of CHD4-bound DEGs have higher expression in *Chd4^−/−^* compared to wild type cells (Fig. 3E, right panel), and there is a statistically significant association between DEGs having a CHD4 peak and being upregulated in *Chd4^−/−^* cells (FET p-value << 0.05, OR = 3.6) (Fig. 3F), suggesting that the main direct effect of CHD4 is repression of transcription.

Altogether, our results indicate that CHD4 has a broad repressive effect on gene expression in differentiating spermatogonia and in cells at early stages of meiosis.

### CHD4 regulates genes involved in spermatogonia differentiation and early meiotic cells

To identify biological processes and pathways directly regulated by CHD4, we performed pathway over-representation analysis on DEGs associated with a CHD4 peak, grouped by cluster and fold change sign (Table S4). Among the pathway databases utilized to test for overrepresentation are GO biological terms (GO:BP) and TRANSFAC^21^, the latter which associates genes with transcription factors according to the motives found at gene’s regulatory regions. In cluster GC_2 (differentiating spermatogonia) and GC_3 (pre-leptotene to zygotene cells), downregulated DEGs were significantly enriched for GO:BP terms such as “spermatogenesis” and *“*spermatid differentiation”. For upregulated DEGs, the terms “chromatin remodeling”*, “*regulation of gene expression”*, “*cell cycle process” and *“*cell differentiation” were enriched in both GC_2 and GC_3 DEGs (Table S4). These findings support a role for CHD4 in regulating key gene expression programs associated with cell differentiation during spermatogenesis. We also observed several transcription factors motives overrepresented in regulatory regions of DEGs from GC_2 and GC_3, such as KAISO (*Zbtb33*), ZF5 (*Zbtb14*), FOXN4 and CTCF (see below), among others.

To specify the role of CHD4 in early germ cell differentiation, we first identified genes involved in developmental transitions, by assessing which genes are differentially expressed between consecutive clusters in wild type cells (GC1→GC2, GC2→GC3, etc.). This analysis showed several known regulators of spermatogonial differentiation, including *Zbtb16* (*a.k.a*. *Plzf)*, *Kit, Gfra1, and Etv5*, as well as early meiotic prophase I markers such as *MeioB*, *Sycp3, Sycp2 and Dmc1 among others* (Table S5). To test whether CHD4 influences these transition-specific gene expression changes, we examined the overlap between these genes and those deregulated in *Chd4^−/−^* cells. Fisher’s exact test revealed an association between *Chd4^−/−^* DEGs, and genes involved in germ cell differentiation across all transitions (GC1→GC2, GC2→GC3, GC3→GC4; Fig. 4A). In all cases, ∼ 40% of genes involved in the transition were also differentially expressed in either cluster in *Chd4^−/−^* cells (Fig. 4A, bar plots). These findings suggest that CHD4 contributes to the regulation of gene expression programs required for the transition from spermatogonia to early prophase I.

**Figure 4.**
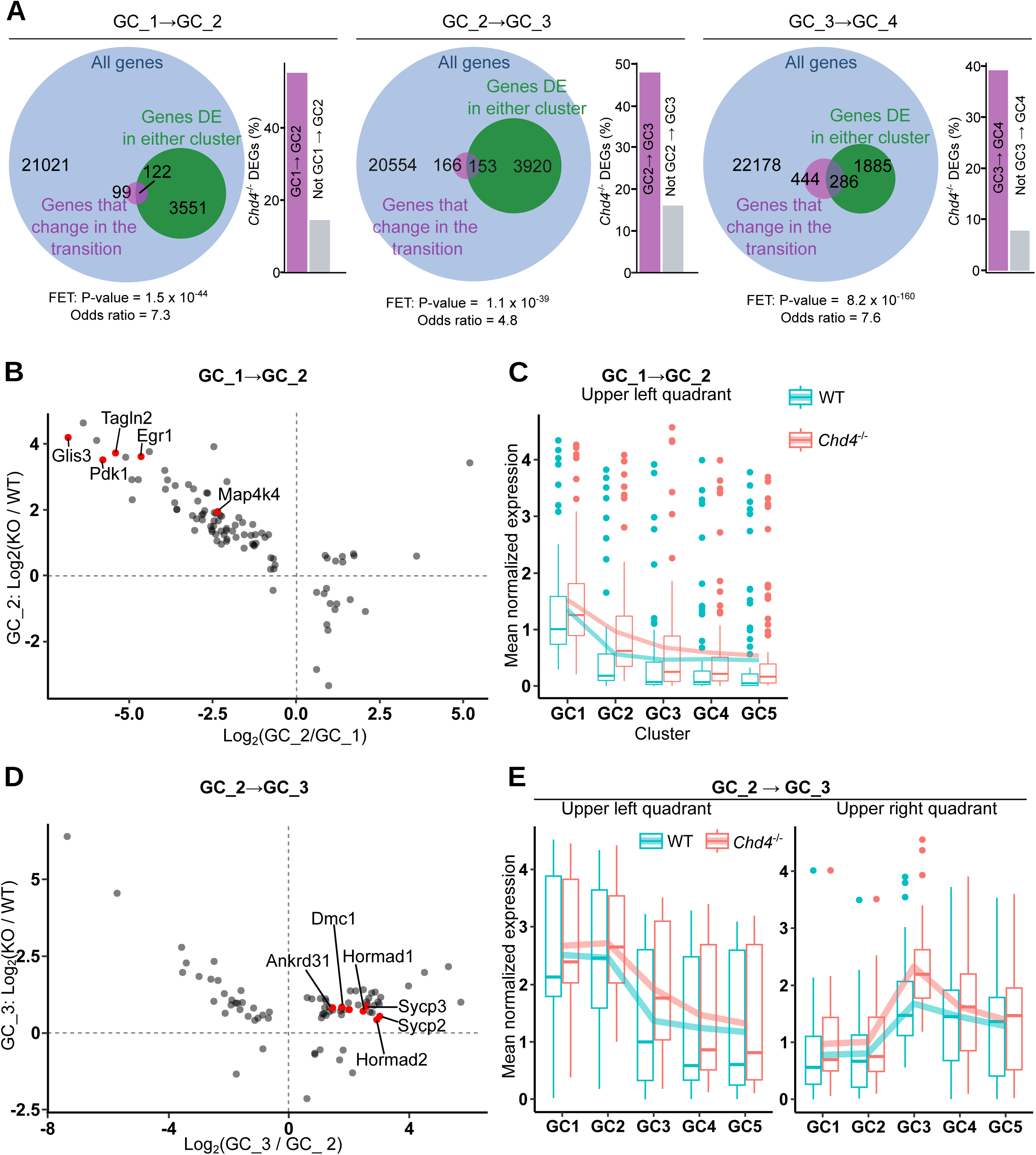
CHD4 modulates transcription of genes involved in spermatogonia differentiation and in transition to meiosis. **(A)** Association between *Chd4^−/−^* DEGs and genes involved in wildtype differentiation. For each transition between two consecutive clusters (GC1→GC2, GC2→GC_3 and GC3→GC4), left: Euler plot depicting the relation between genes that change expression between clusters in Chd4wildtype cells (*i.e.* genes that change during differentiation), and genes that are differentially expressed in *Chd4^−/−^* samples in either of the two clusters. Right: percentage of *Chd4^−/−^* DEGs (from either clusters involved in the transition) that change expression during the transition in wildtype cells (’GCn→GCn+1’), or that do not change expression during the transition in wildtype cells (’Not GCn→GCn+1’). p-value and odds ratio for Fisher’s exact test on categories ‘gene is differentially expressed in either cluster in *Chd4^−/−^* and gene is differentially expressed between those two clusters in wild type cells’. **(B)** Scatter plot depicting the change in expression in wild type cells in the transition from GC_1 to GC_2 in the x-axis (log_2_(GC_2/GC_1)), and the change in expression between *Chd4^−/−^*and wild type cells from cluster GC_2 in the Y-axis (log2(*Chd4^−/−^*/WT)). Each dot represents a gene and only genes annotated with a Chd4 peak and a statistically significant change in expression in both conditions are shown (FDR < 0.05). **(C)** Box plots depicting mean normalized expression per cluster and genotype for the genes found in the ‘upper left quadrant’ in panel B, *i.e.* genes bound by CHD4 that reduce their expression upon transition from cluster GC_1 to GC_2 in wild type cells and that have a higher expression in *Chd4^−/−^* mice with respect to wild type in cluster GC_2. Line represents the mean of the values in the box plot. **(D)** Scatter plot depicting the change in expression in wild type cells in the transition from GC_2 to GC_3 in the x-axis (log_2_(GC3/GC2)), and the change in expression between Chd4^-/-^ and wild type cells from cluster GC_3 in the Y-axis (log_2_(*Chd4^−/−^*/WT)). Each dot represents a gene and only genes associated with a Chd4 peak and a statistically significant change in expression in both conditions are shown (adjusted p-value < 0.05). **(E)** The box plots depict mean normalized expression per cluster and genotype for the genes found in the ‘upper left quadrant’ or ‘upper right quadrant’ in panel D. Line represents the mean of the values in the box plots.

To assess whether CHD4 directly regulates genes involved in differentiation, we repeated the analysis considering only *Chd4^−/−^* DEGs annotated with a CHD4 peak. Since CHD4-peaks lists are from spermatogonia or preleptotene/leptotene, we limited this analysis to clusters GC_1 to GC_3. For both the GC_1→GC_2 and GC_2→GC_3 transitions, *Chd4^−/−^* DEGs annotated with a CHD4 peak were associated with genes involved in differentiation (Fig. S5). Again, this supports a model in which CHD4 directly regulates gene expression programs governing spermatogonia differentiation and entry into meiosis.

We next examined how transcriptomic changes during the GC_1→GC_2 transition in wild type cells correlate with gene expression differences between *Chd4^−/−^* and wild type cells within GC_2 (Fig. 4B). For this analysis, we focused on genes that meet the following conditions. They are a-transcriptionally regulated during spermatogonia (i.e., differentially expressed in GC_1→GC_2 developmental transition in wild type cells) and b-directly regulated by CHD4 (i.e., have a peak of CHD4 and be differentially expressed in *Chd4*^−/−^ in GC_2). We observed that most of these genes are downregulated upon spermatogonia differentiation in wild type mice and are upregulated in *Chd4^−/−^* cells. These findings suggest a role for CHD4 in promoting downregulation of transcripts upon spermatogonia differentiation.

To visualize the transcriptional changes of the genes described above in the context of development, we plotted their average expression across clusters, in wild type and *Chd4^−/−^* samples (Fig. 4C and S6A). In wild type cells, these genes are highly expressed in undifferentiated spermatogonia (GC_1), become downregulated in GC_2, and remain low in subsequent stages (Fig. 4B-C and S6A). In contrast, *Chd4^−/−^* cells retain elevated expression of many of these genes in GC_2, with downregulation delayed until later stages. These results indicate that CHD4 plays a critical role in repressing transcription during spermatogonia differentiation, particularly by ensuring timely silencing of genes associated with the undifferentiated state.

A more complex pattern emerged when analyzing the transition from GC_2 to GC_3, which marks meiotic entry. Again, we focused on genes that meet the following conditions. They are a-transcriptionally regulated during meiotic entry (i.e., differentially expressed in GC_2→GC_3 developmental transition in wild type cells) and b-directly regulated by CHD4 (i.e., have a peak of CHD4 and be differentially expressed in *Chd4^−/−^* in GC_3). Most of these genes are upregulated in *Chd4^−/−^* cells in GC_3 (Fig. 4D, upper quadrants), indicating an intrinsic repressive function of CHD4. Some of these genes are downregulated during GC_2→GC_3 transition while others are upregulated (Fig. 4, upper left and right quadrant, respectively). The downregulated genes are highly expressed in wild type spermatogonia (GC_2 and/or GC_1) and become downregulated at the onset of meiosis (GC_3) and remain with low expression thereafter (Fig. 4D and E, upper left quadrant, and Fig. S6B). In contrast, in *Chd4^−/−^* cells we observe that these genes have a more gradual decline in expression. Several of these genes (Fig. 4D, upper left quadrant) are involved in ribosomal function (Table S6). These findings suggest that CHD4 plays a critical role in repressing spermatogonial gene expression during the transition into meiosis.

As for the set of genes directly regulated by CHD4 that are upregulated during the transition (Fig. 4D, upper right quadrant), we observe that in *Chd4^−/−^* cells, their expression is markedly elevated in GC_3 but have proper expression in subsequent stages (Fig. S6C). These findings suggest that CHD4 contributes to the regulation of early meiotic gene expression by tempering transcriptional activation at meiotic entry.

Altogether, our results support a model in which CHD4 is required for proper expression of genes important for spermatogonia development, meiotic onset, and early meiosis.

### CHD4 and CTCF overlap at active promoters and enhancers in meiotic cells

We observed motives associated to specific transcription factors overrepresented in regulatory regions of DEGs with a peak of CHD4. Among these, we found CTCF. Given the proposed prominent role for CTCF in transcriptional regulation in the male germline, possibly directing early gametogenic transcriptional developmental programs and transitions^22^, we further investigated whether CTCF role is connected to CHD4. To do so, we performed CUT&RUN experiments using WIN18,446/retinoic acid-synchronized testes to obtain enriched fractions of preleptotene/leptotene spermatocytes. We observed 22,160 CTCF consensus peaks in wild type spermatocytes and 33,911 consensus peaks in *Chd4^−/−^* spermatocytes (Fig. S7A-C). In wild type preleptotene/leptotene cells, CTCF overlaps with nearly half of CHD4 binding sites (Fig. 5A). We characterized CTCF and CHD4 common binding sites according to the distance to the closest TSS. Sites were classified as promoters if their distance was between - 1,000 bp and +300bp; as distal, if their distance was below -2,000 bp or above +1,000 bp; or intermediate, if the distance fell between these values. Fig. 5B shows that CTCF binding sites that overlap with CHD4 are more likely to be on the promoter region compared to those that do not overlap with a CHD4 binding site. In a similar fashion, CHD4 more likely overlaps with CTCF at promoters (Fig. 5C).

**Figure 5.**
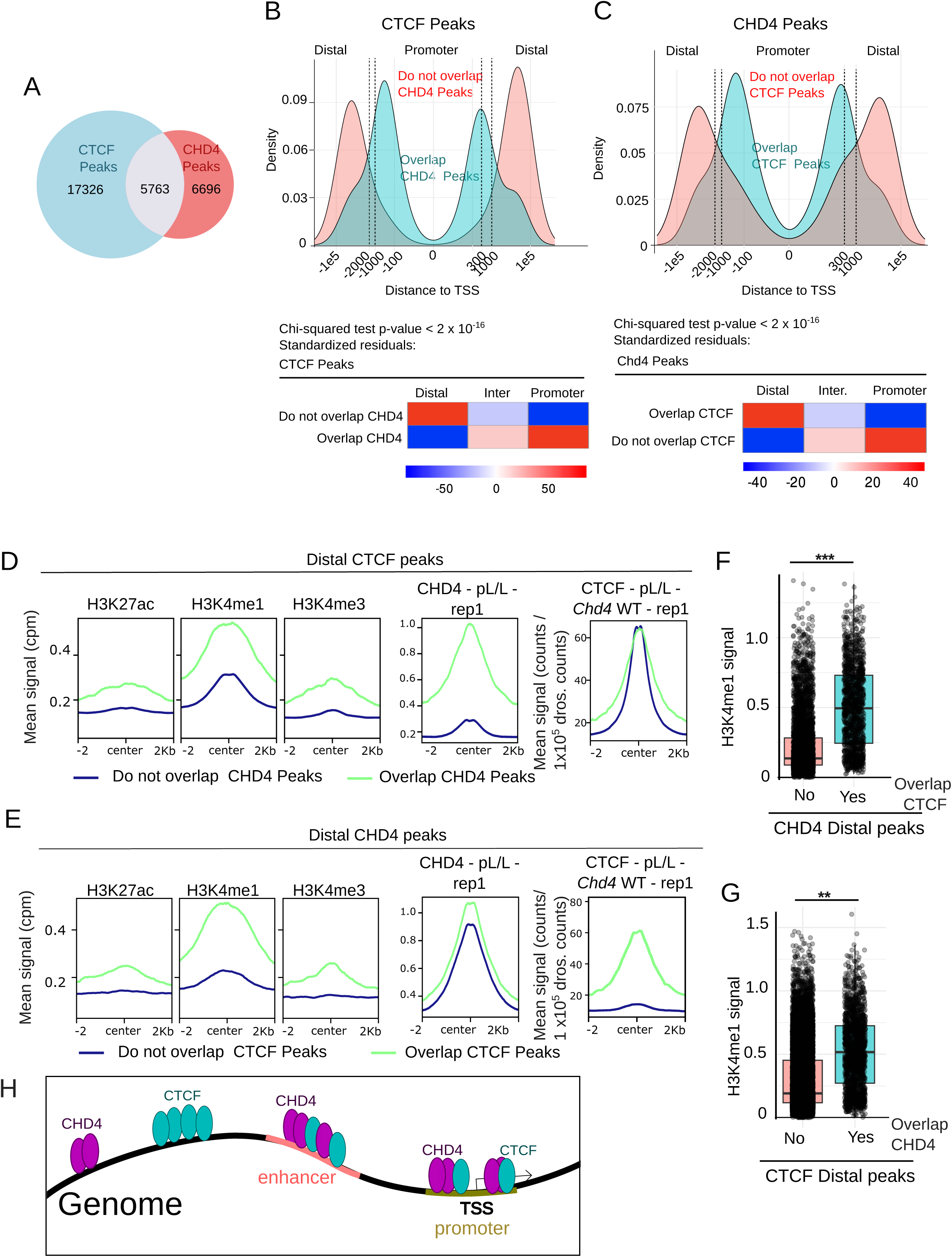
CHD4 overlaps CTCF at active promoters and enhancers. **(A)** Euler plot depicting CTCF and CHD4 peaks overlap at preleptotene. **(B-C**) Top: Distribution of distances of CTCF (B) and CHD4 (C) preleptotene consensus peaks to the closest TSS, grouped by whether they overlap a CHD4 (B) or CTCF(C) peak (blue) or not (red). Vertical dotted lines mark −2000, −1000, +300, and +1000 bp relative to the TSS. Peaks were classified as promoter (−1000 to +300 bp), distal (< −2000 or > +1000 bp), or intermediate (between −2000 to −1000 bp or +300 to +1000 bp). Bottom: results of a chi-squared test assessing the association between TSS distance category and overlap with the other protein. Shown are p-values and standardized residuals, where positive residuals indicate enrichment and negative residuals indicate depletion relative to expectation. **(D)** Mean signal profiles of histone marks H3K27ac, H3K4me1 and H3K4me3, CHD4 (replicate 1) and CTCF (replicate 1) in wild type preleptotene cells. Profiles are centered at distal CTCF preleptotene consensus peaks that either overlap (green) or do not overlap a CHD4 preleptotene consensus peak. **(E)** Mean signal of same proteins as in D but centered at distal CHD4 preleptotene consensus peaks that either overlap (green) or do not overlap a CTCF preleptotene consensus peak. **(F-G)** H3K4me1 signal at distal CTCF (F) or CHD4 (G) preleptotene consensus peaks, grouped by overlap with CHD4(F) or CTCF (G) preleptotene consensus peaks (Yes/No). Each point represents an individual peak; boxplots show median and interquartile range. Differences between groups were tested using the Wilcoxon rank-sum test (p-value < 2×10⁻^16^ in both cases). Effect sizes were measured with Cliff’s delta (F: 0.583, large effect size, ***; G: 0.449, medium, **). **(H)** Cartoon depicting the proposed model: in preleptotene, CHD4 overlaps CTCF peaks at enhancers (distal peaks, high H3K4me1 signals) and promoters (proximal peaks, high H3K4me3, H3K27ac). Distal non overlapping peaks have on average, little signal for all tested histone marks.

We then tested the association of CTCF and CHD4 common binding sites with histone post-translational modifications usually associated with different genomic features and/or transcriptional states^23^. CTCF and CHD4 promoter binding sites show marks for active promoters (H3K4me3 and H3K27ac) and absence of repressive marks (H3K9me3 and H3K27me3) (Fig. S8 and Fig. S9), indicating that both proteins overlap mostly at active promoters. In the case of distal peaks, overlapping CHD4 and CTCF had signal of H3K4me1, indicating that CTCF and CHD4 also overlap at enhancers. Both proteins have additional peaks that do not overlap and tend to be distal to TSSs (Fig. 5 B and C) and with reduced enhancer histone marks (Fig. 5 D-G).

We conclude that CHD4 and CTCF may work together in active promoters and enhancers (Fig. 5H).

### CHD4 regulates gene transcription by regulating CTCF binding at promoters and enhancers

To assess whether CHD4 influences CTCF binding, we analyzed CUT&RUN CTCF signal in *Chd4^−/−^* versus wild type control preleptotene/leptotene spermatocytes. The results showed a broad increase in CTCF binding to chromatin in the absence of CHD4. More than 19,000 regions displayed higher CTCF signal in *Chd4^−/−^* mice (positive differentially bound regions, DBRs), whereas only about 1,000 regions showed reduced signal (negative DBRs) (FDR ≤ 0.05; Fig. 6A).

**Figure 6.**
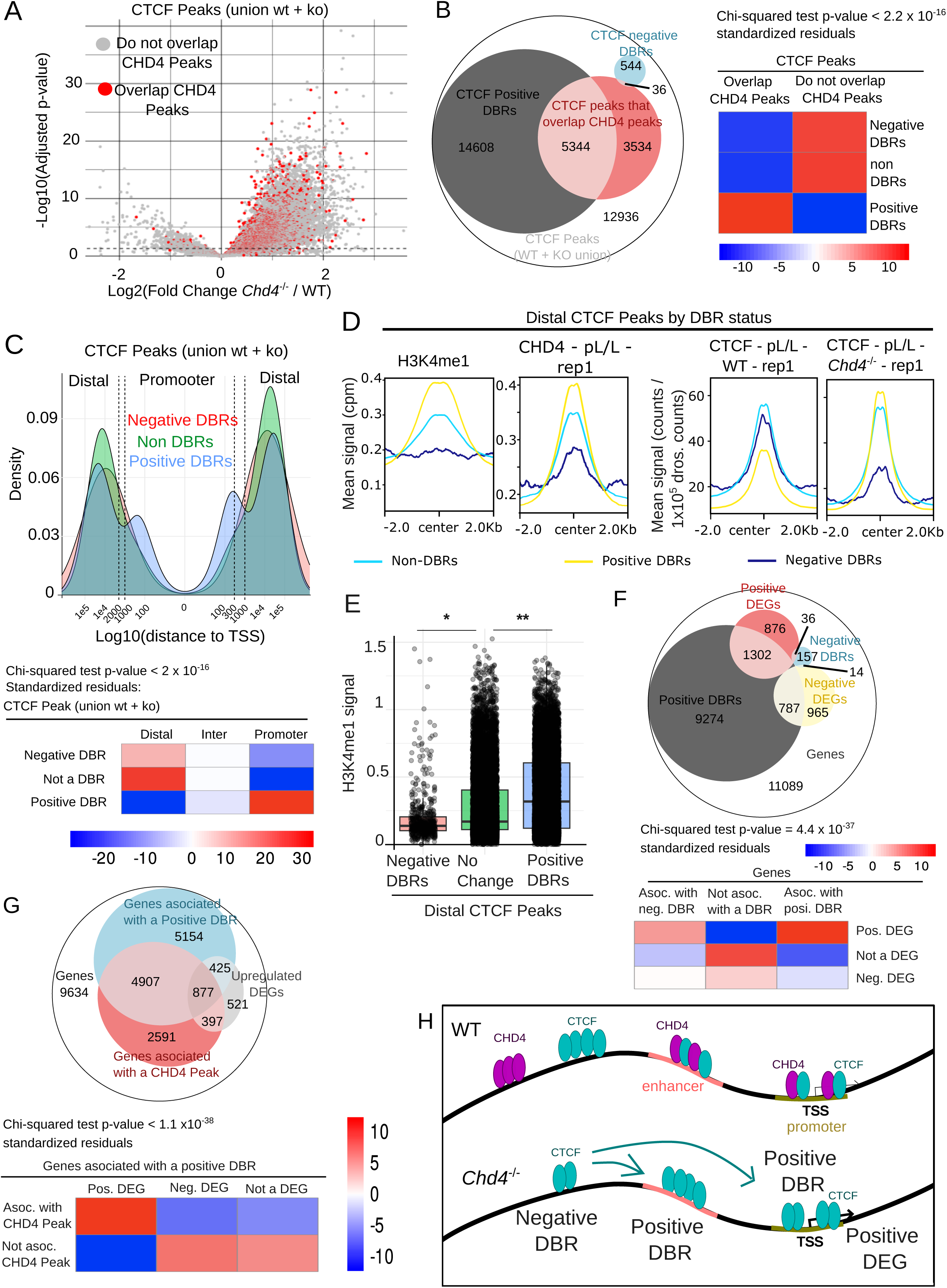
CHD4 regulates transcription in preleptotene by inhibiting CTCF binding at active promoters and enhancers. **(A)** Volcano plot of CTCF differential binding analysis results. The analysis was performed on the union of CTCF consensus peaks in preleptotene from *Chd4^−/−^* and wild type mice. CTCF binding regions overlapping a CHD4 peak are highlighted in red. The dotted horizontal line indicates the significance threshold (FDR < 0.05). **(B)** Left: Euler plot showing the number of CTCF regions with statistically significant differentially bound regions (DBRs) in *Chd4^−/−^* with respect to wild type mice. Regions with higher CTCF binding in the *Chd4^−/−^* mice are labeled as positive and regions with lower binding as negative. Right: Chi-squared test results (p-value and standardized residuals in color scale) for association between CTCF binding regions overlapping a CHD4 peak and their DBR status (negative, positive or non-DBR). Positive standardized residuals indicate an enrichment of observed over expected counts, whereas negative residuals indicate a depletion. **(C)** Top: Distribution of distances of CTCF preleptotene binding regions (formed by union of CTCF consensus peaks in preleptotene at *Chd4^−/−^* and wild type mice) to the closest TSS, grouped by their DBR status. Dotted lines mark −2000, −1000, +300, and +1000 bp relative to the TSS. Peaks were classified as promoter (−1000 to +300 bp), distal (< −2000 or > +1000 bp), or intermediate (between −2000 to −1000 bp or +300 to +1000 bp). Below: Chi-squared test results (p-value and standardized residuals in color scale) for the association between distance to TSS (columns) and DBR status (rows). **(D)** Mean signal profiles of H3K4me1, CHD4 (replicate 1) and CTCF (replicate 1) in wild type and *Chd4^−/−^* (replicate 1) preleptotene cells. Profiles are centered at distal CTCF preleptotene binding regions (union of consensus peaks in Chd4-wildtype and *Chd4*^-/-^ mice), grouped by DBR status (positive, negative or non-DBRs). **(E)** H3K4me1 signal at distal CTCF preleptotene binding regions (union of consensus peaks in *Chd4* wild type and *Chd4^−/−^* mice), grouped by DBR status. Each point represents an individual peak; boxplots show median and interquartile range. Differences across groups were assessed with a Kruskal–Wallis test (*p* < 2.2 e-16), followed by post-hoc pairwise Wilcoxon rank-sum tests. p-values of the pairwise comparisons indicated above the barplots (* = 1.8 x 10^-14^; ** < 2.2 x 10^-16^). **(F)** Top: Euler plot showing overlap between *Chd4^−/−^* positive and negative DEGs among GC_2 and GC_3 clusters, and genes associated to at least one positive or negative DBR (’Positive DBRs’ and ‘Negative DBRs’). The following genes and the DBRs associated to them were excluded from the analysis: genes that had different fold change sign in GC_2 and GC_3, genes annotated to both a positive and a negative DBR, DBRs associated to genes that were not detected in the scRNA-seq full dataset. In total, 318 genes and 4340 DBRs were excluded. Bottom: Chi-squared test p-value and standardized residuals for genes categorized according to DBR and DEG status. (G) Top: Euler plot showing association between positive DEGs, genes associated to a positive DBR and genes associated with a CHD4 peak. Same genes and DBRs as in (F) were excluded from analysis. Below: Chi-squared test p-value and standardized residuals (color scale) for positive DBRs categorized according to DEG status and whether or not they overlap a CHD4 peak. **(H)** Model: CHD4 represses transcription of a subset of genes by binding to their enhancers and/or promoters and preventing CTCF binding at these regulatory sites. In the absence of CHD4, CTCF binding increases at promoters and enhancers, which in turn leads to a depletion of CTCF at distal sites not associated with these gene regulatory elements.

When intersecting CTCF DBRs with CHD4 binding sites at the preleptotene/leptotene stage, we found a statistically significant association between CTCF DBR status and CHD4 peak overlap (Fig. 6A and B). Positive DBRs (higher CTCF signal in *Chd4^−/−^*) were enriched among CTCF binding sites overlapping CHD4, whereas negative DBRs were enriched among those lacking CHD4 overlap (Fig. 6B). These results are consistent with CHD4 acting directly at CTCF binding sites to limit CTCF occupancy.

Next, we analyzed the distribution of CTCF peak distances to the nearest TSS (classified as promoters, intermediate, or distal) in relation to their DBR status. We found that CTCF binding sites at promoters were enriched in positive DBRs, whereas distal CTCF binding sites were enriched in non-DBRs (Fig. 6C).

We also analyzed the signal of histone marks at CTCF sites in wild type and *Chd4^−/−^* mice, grouping sites by their distance to the nearest TSS and by their DBR status. Proximal CTCF binding sites show similar levels of active promoter-associated marks (H3K4me3 and H3K27ac) regardless of DBR status (Fig. S10). Distal CTCF sites, however, displayed DBR-dependent differences in H3K4me1 signal (enhancers). This is, positive DBRs had the highest levels of H3K4me1, followed by non-DBRs, and finally negative DBRs, the latter showing little to no signal (Fig. 6D, E). Together, these results suggest that CHD4 regulates CTCF binding both at active promoters and enhancers.

Negative DBRs are mostly distal (Fig. 6C), do not overlap with CHD4 peaks (Fig. 6B) and lack H3K4me1 marks (Fig. 6D), indicating that they are not enhancers. The decreased binding of CTCF to this type of region could be explained by the increased binding of CTCF to other regions, which depletes a limited pool of this protein (Fig. 6H).

Finally, we analyzed whether CTCF differential binding was associated with transcriptional changes in *Chd4^−/−^* spermatocytes. Out of 2,238 upregulated DEGs in clusters GC_2 or GC_3 (differentiating spermatogonia and preleptotene/leptotene to zygotene, respectively) 1,302 (58%) were associated with a positive DBR—a statistically significant association as determined by a chi-squared test (Fig. 6F). Moreover, among positive DBRs, those that overlapped a CHD4 peak were preferentially associated with upregulated DEGs (Fig. 6G). Taken together, these results suggest that CHD4 represses transcription, in part, by reducing CTCF binding at active promoters and enhancers (Fig. 6H).

## Discussion

Sexual reproduction in mammals depends on the sustained generation of haploid gametes, a process governed by tightly regulated developmental stages and transitions including spermatogonia maintenance and differentiation, meiosis, and spermiogenesis. These transitions are orchestrated by chromatin remodeling complexes and architectural proteins such as CTCF, which together shape the transcriptional landscape necessary for germ cell maturation. While both CHD4 and CTCF have been individually implicated in early spermatogonial development, their mechanistic interplay and role during later stages of spermatogenesis remained unclear. Here, we demonstrate that by acting on key developmental gene expression programs CHD4 plays a dual role in gamete development: it is important for normal spermatogonial development and also for progression of meiosis. Through integrative analyses of genome-wide chromatin binding and single-cell transcriptomics, we uncover that CHD4 regulates and directly modulates CTCF chromatin binding and transcriptional function in developing gametes. This study provides new insights into the action of CHD4 and CTCF on spermatogonia and meiotic cells and uncover a previously unrecognized CHD4–CTCF regulatory axis that coordinates early testis development and meiotic progression, ensuring faithful formation of haploid gametes and accurate genome transmission to the next generation.

We determined that spermatocyte development arrest occurs at mid stages of meiosis I prophase with an apparent moderate effect on DNA repair and severe defects and homologous chromosome synapsis. Deficient repair of double strand breaks in *Chd4^−/−^* spermatocytes is not surprising as CHD4 has been directly implicated in the response to damaged DNA in somatic cells models. For example, CHD4 is recruited to artificially generated DNA damage sites and affects DNA repair through different pathway^16,17^. CHD4 also interacts with MCPH1, and its loss interferes with the recruitment of MCPH1 to damage sites^24^. Importantly, MCPH1 regulates the recruitment of RPA, RAD51 and BRCA2 to DSB sites undergoing recombination repair. Given its positive impact on survival following irradiation, current models predict that CHD4 in somatic cell models promotes a permissive chromatin environment for proper DSB repair^16,17^, which may explain the phenotype we observed in meiotic cells. However, analysis of RPA, RAD51, and MLH1 foci formation in *Chd4^−/−^* spermatocytes revealed no difference with respect to wild type, suggesting cell type-specific differences, with a reduced effect of CHD4 on meiotic recombination DNA repair. *Chd4^−/−^* spermatocytes show incomplete homologous chromosome synapsis. Evaluated by SYCP3 and SYCP1 immunostaining, the most advanced spermatocytes showed fully developed axes, but the homologous pairs remained incompletely synapsed at the centromeric chromosome ends. In these chromosome areas, the axial element of the synaptonemal complex exhibits abnormal deposition of SYCP3. Abnormal DNA repair may be related to defects in chromosome synapsis observed in *Chd4^−/−^* spermatocytes. Alternatively, we observed a transcriptional effect of CHD4 on meiotic-specific genes participating in formation and/or dynamics of the synaptonemal complex (e.g., *Hormad*, *Sycp2*, *Sycp3*), which could explain the observed phenotype. Together, our mutant analysis suggests an essential role for the CHD4 ATPase in normal mammalian male meiotic progression. However, because of the possible vast spectrum of CHD4 functions, it is difficult to solely use phenotypic characterization to pinpoint the specific molecular causes of failure.

To gain mechanistic insights, we combined analysis of genome wide chromatin binding with its functional implications in cell specific gene transcription. We uncovered previously unappreciated regulatory functions for CHD4 and CTCF in spermatogonia development, meiotic commitment, and meiotic progression. In spermatogonia, we observe that CHD4 is a key driver of the transcriptional repression necessary for spermatogonia to differentiate and to enter meiosis. This analysis showed several known genes directly involved in spermatogonial maintenance and differentiation, including *Kit*^25^, *Dmrt1*^26^, *Stra8*^27^, *Foxo1*^28^, and *Tagln2*^29^ among others. In previous work, using Ddx4-cre, CHD4 was seeing to promote the survival of spermatogonia stem cells by promoting the expression of self-renewal genes and inhibiting the expression of genes characteristic of more differentiated stages^8,9^, or directly inhibiting expression of pro-apoptotic genes^11,30^. The differences observed in gene targets and pathways affected by CHD4 could be ascribed to deletion of *Chd4* in different spermatogonia stages and reflects the plasticity of CHD4 to perform different roles in different cellular contexts, even within the same cell linage.

The transcriptional control effect of CHD4 is extended to genes participating in meiosis specific processes such as recombination-mediated repair of double strand breaks (*Meiob*, *Dmc1*, *Ankrd31*) and homologous chromosome pairing and synapsis (*Hormad1*, *Hormad2*, *Sycp2*, *Sycp3*).

Our data provides a substantial conceptual advance by integrating CTCF and CHD4 regulatory functions. Because most CTCF binding sites are located at nucleosome-free regions, its binding is likely regulated by chromatin remodeling. Indeed, we show that in certain developmental contexts, such as in spermatogonia and early meiosis, CHD4 chromatin remodeling works as a pervasive regulator of CTCF chromatin binding. In this model, CHD4 control of chromatin accessibility prevents inappropriate binding of CTCF at selected groups of promoters and enhancers. This CHD4 action, may explain its role as a paramount repressive factor preventing inappropriate expression of genes, ultimately required for key spermatogenic developmental stages and transitions.

Together, our data adds a novel layer of understanding in how chromatin remodelers integrate high order chromatin structure and transcriptional outcomes by influencing the function of central regulatory factors such as CTCF.

## Material and Methods

### Mice

*CHD4*-floxed mice (*CHD4^fl/fl^)* have been described^9^. Transgenic *Cre* recombinase mice Ngn3-Cre**^FVB^**(Cg)**^-Tg^**(Neurog3–cre)**^C1Able/J^** was purchased from The Jackson Laboratory (Bar Harbor, ME). *Chd4* gonad-specific knockouts and wild type heterozygotes littermates were obtained from crosses between female homozygous *flox/flox* mice with male heterozygous Cre/+; *Chd4* Wt/flox mice.

All experiments conformed to relevant regulatory standards guidelines and were approved by the Oklahoma Medical Research Foundation-IACUC (Institutional Animal Care and Use Committee).

### Mice Genotyping

Characterization of wild type and *floxed* alleles was carried out by PCR using the following oligonucletides: *Chd4* forward 5’-TCC AGA AGA AGA CGG CAG AT and *Chd4* reverse 5’-CTG GTC ATA GGG CAG GTC TC (Floxed ∼400 bp, Wt ∼290 bp). Ngn3-Cre detection was performed using the following primer set: mutant Reverse 5’-ACA TGT CCA TCA GGT TCT TGC, common forward 5’-GGC ACA CAC ACA CCT TCC A, wild type Reverse 5’-AGT CAC CCA CTT CTG CTT CG (Transgene = ∼165 bp, Wt = 197 bp).

### Western blot cell lysates

Total testis or enriched cells fractions were lysed in ice-cold protein extraction buffer containing 0.1 % Nonidet P-40, 50 mM Tris-HCl, pH 7.9, 150 mM NaCl, 3 mM MgCl2, 3 mM EDTA, 10 % glycerol, 1 mM DTT, 1 mM PMSF and protease inhibitors (ThermoFisher Scientific, A32965) followed by sonication (3 pulses of 10 seconds) using micro ultrasonic cell disrupter (Kontes). The relative amount of protein was determined measuring absorbance at 260 nm using NanoDrop 2000c spectrophotometer (ThermoFisher Scientific). Proteins were solubilized with 2 X sample buffer (4 % SDS, 160 mM Tris-HCl, pH 6.8, 20 % glycerol, 4 % mM β-mercaptoethanol, and 0.005 % bromphenol blue) and 30 µg/lane of sample were separated by 4–15 % gradient Tris-acetate SDS-PAGE and electro transferred to PVDF membrane (Santa Cruz Biotechnology, sc-3723). The blots were probed with individual primary antibodies and then incubated with HRP-conjugated goat anti-mouse or rabbit antibody as required. In all blots, proteins were visualized by enhanced chemiluminescence and images acquired using Western Blot Imaging System c600 (Azure Biosystems). ImageJ software was used for quantification of non-saturated bands and tubulin was used for normalization. The antibodies used are detailed in Table S7.

### Histology and immunostaining

Testes and ovaries were dissected, fixed in 10 % neutral-buffered formalin (Sigma) and processed for paraffin embedding. Serial sections from paraffin wax-embedded testes (5–8-μm) or ovaries were positioned on microscope slides and analyzed using either Hematoxylin and Eosin staining or a TUNEL assay (Roche). For immunostaining analysis, tissue sections were deparaffinized, rehydrated and antigens were recovered in sodium citrate buffer (10 mM Sodium citrate, 0.05 % Tween 20, pH 6.0) by heat/pressure-induced epitope retrieval. Incubations with primary antibodies were carried out for 12 h at 4 °C in PBS/BSA 3 %. Primary antibodies used in this study are detailed in Table S7. Following three washes in 1 X PBS, slides were incubated for 1 h at room temperature with secondary antibodies. A combination of fluorescein isothiocyanate (FITC)-conjugated goat anti-rabbit IgG (Jackson laboratories) with Rhodamine-conjugated goat anti-mouse IgG and Cy5-conjugated goat anti-human IgG each diluted 1∶450 were used for simultaneous triple immunolabeling. Slides were subsequently counterstained for 3 min with 2 µg/ml DAPI containing Vectashield mounting solution (Vector Laboratories) and sealed with nail varnish. We use Zen Blue (Carl Zeiss, inc.) for imaging acquisition and processing.

### Cytology

We employed established experimental approaches for the visualization of chromosomes in chromosome surface spreads [18]. Incubations with primary antibodies (Table S7) were carried out for 12 h at 4 °C in 1 X PBS plus BSA 2 %. Following three washes in 1 X PBS, slides were incubated for 1 h at room temperature with secondary antibodies. A combination of fluorescein isothiocyanate (FITC)-conjugated goat anti-rabbit IgG (Jackson laboratories) with Rhodamine-conjugated goat anti-mouse IgG and Cy5-conjugated goat anti-human IgG each diluted 1∶350 was used for simultaneous immunolabeling if required. Slides were subsequently counterstained for 3 min with 2 μg/ml DAPI containing Vectashield mounting solution (Vector Laboratories) and sealed with nail varnish. We used Zen Blue (Carl Zeiss, Inc.) for imaging acquisition and processing.

### Primary spermatocyte enrichment

Synchronized spermatocytes from the first spermatogenic wave were obtained as described in^31^. Briefly, 2 dpp mice were orally gavaged for 7 consecutive days with WIN 18,446 (dissolved in 0.5% aqueous tragacanth suspension) to arrest germ cells as spermatogonia. After one day without treatment, (10 dpp), mice were injected with retinoic acid (RA) to induce their coordinated maturation. Testes were collected at 7 days after RA injection corresponding to cell populations enriched in preleptotene/leptotene cells.

Cells obtained from a single-cell suspension protocol were washed in DMEM containing 5% FBS and centrifuged for 15 min at 150 × g, room temperature. The cell pellet was washed once in DMEM and resuspended in DMEM supplemented with 10% FBS and 1% antibiotic–antimycotic solution. Cells were plated onto cell culture plates and maintained in a humidified incubator at 37 °C with 5% CO₂ for 4 h. After incubation, the non-adherent (floating) germ cells were collected for further processing.

Purity and spermatogenic stage of cells were assessed by immunofluorescence of chromosome spreads using γH2AX and SYCP3 antibodies.

### Generation of single cell suspension

Testis from mice with the indicated age were removed and placed into a Petri dish containing RT Dulbecco’s Modified Eagle Medium. After detunication, slightly stretched seminiferous tubules were placed in fresh RT DMEM medium supplemented with collagenase at a final concentration of 0.45 mg/ml under gentile agitation for 10 min. After washing with DMEM, the tubules were treated with DNAse I (25 U, 2 min) and then with Trypsin (0.05% w/v) and incubated for 30 min at room temperature. The reaction was stopped with 5% FBS followed by 10 min of incubation at room temperature. Single cell suspension was obtained by pipetting followed by filtration through a 40-µm-poresize cell strainer.

### CUT & RUN

#### Cell preparation and genomic DNA purification

Cells were harvested by centrifugation (300 × g for 15 min at 4°C) and washed three times in washing buffer (20 mM HEPES pH 7.5, 150 mM NaCl, 0.5 mM spermidine). After the final wash, they were resuspended in 1 ml of the same buffer. An appropriate number of cells (500,000 for CTCF and 350,000 for CHD4) was added to concanavalin A-coated beads, pre-washed and resuspended in binding buffer (20 mM HEPES, 10 mM KCl, 1 mM CaCl2, 1 mM MnCl2), and incubated for 10 minutes at room temperature. Cells attached to beads were resuspended in antibody buffer (20 mM HEPES pH 7.5, 150 mM NaCl, 0.5 mM spermidine, 2 mM EDTA, 0.1% (v/v) digitonin, 1 mM PMSF, 10 mM β-glycerophosphate and 1× Protease Inhibitors Cocktail) containing the appropriate primary antibody at a 1:100 dilution. Samples were incubated on a mixing platform at 4°C overnight. Next day, the beads were washed three times in Dig-wash buffer (20 mM HEPES pH 7.5, 150 mM NaCl, 0.5 mM spermidine, 0.05% (v/v) digitonin, 1 mM PMSF, 10 mM β-glycerophosphate, 1× protease inhibitor cocktail) and then resuspended in 50 μl of the same buffer containing pA-MNase at a final concentration of 700 ng/ml and incubated on a mixing platform for 1 hour. After incubation, the beads were washed again three times in Dig-wash buffer and resuspended in 100 μl of this solution. pA-MNase was activated by adding CaCl2 to a final concentration of 2 mM and incubated on ice for 30 minutes. The reaction was stopped by the addition of STOP buffer (340 mM NaCl, 20 mM EDTA, 4 mM EGTA, 0.05% (v/v) digitonin, 0.1 mg/ml RNase A, 200 pg/ml Drosophila spike-in DNA). To release DNA fragments from insoluble nuclear chromatin, the samples were incubated for 30 minutes at 37°C. The supernatant was collected and subjected to DNA isolation using a Qiagen kit (REF: 28006) according to the manufacturer’s instructions.

#### CUT & RUN Genomic library preparation

Libraries were prepared using Integrated DNA Technologies (IDT) xGen™ ssDNA & Low-Input DNA Library Prep Kit (cat. # 10009859) in conjunction with IDT xGen UDI Primers Plate 2-8 (cat. # 10009816), following manufacturer’s instructions, and sequenced as paired end 150 bp on an Illumina Novaseq 6000 instrument.

#### Alignment, filtering and generation of coverage track files (bigwig)

For samples prepared using adaptase technology (see table S3), reads 1 and 2 were trimmed 10 bases at the beginning using fastp [36] (version 0.23.2), as recommended by the library preparation kit manufacturer. Further processing was done in parallel for all samples (regardless of the technology used to prepare the library). Reads were adapter-trimmed and quality-pruned using fastp with default settings. Then, reads were aligned to mm10 genome using BWA–MEM (citation) (version 0.7.15) with default settings except for option ‘-M’ for Picard compatibility. For samples with drosophila spike-in DNA, alignment was done to a fusion of drosophila’s d6 and mouse mm10 reference genomes, and next reads aligned to either genome were separated in two separate bam files. From there on, experiments with or without drosophila were processed in parallel. Picard (version 2.21.2, http://broadinstitute.github.io/picard/) and SAMtools (version 1.14) (citation) were used to obtain mapping quality metrics, remove duplicates and filter reads. Only primary alignment reads, not duplicated, properly paired, with a MAPQ > 30, and not placed in mitochondrial chromosome or in unassigned/unplaced sequences were kept. Coverage track files were generated using deepTools (v 3.4.3, citation) bamCoverage with the following arguments: -bs 4, --smoothLength 1, --normalizeUsing CPM. To generate the drosophila normalized converage bigwig files, instead of using -- normalizeUsing CPM argument, --scaleFactor 100,000/total_drosophila_counts was used.

#### Peak calling, consensus peaks generation and annotation

Peaks were called using MACS2 (version 2.2.7.1) using broad mode, an input file, extension 300 for single end experiments, and default values for the rest of the parameters. Peaks were filtered by ENCODE blacklist regions (https://doi.org/10.1038/s41598-019-45839-z).

Consensus peaks for a set of samples constitute all the regions, longer than 10 bp, in which 2 or more peaks overlapped. These were generated using a combination of bedtools multiinter, awk, and bedtools merge.

In the case of spermatogonia, we first generated consensus peak for data obtained from Li *et al.*, 2010. We used that list of consensus peaks as one sample, to finally generate a 2 out of 3 consensus list.

To annotate any set of regions (either consensus peaks or DBRs) to the nearest transcription start site we used HOMER’s annotatePeaks.pl script (https://pubmed.ncbi.nlm.nih.gov/20513432/, version 4.11) together with cell ranger’s gft file.

### Aggregate profiles

To generate aggregate profiles, we used deepTools computeMatrix (arguments reference-point and --skipZeros) and plotHeatmap.

### Genome wide binding correlation analysis

To plot the correlation between genome wide binding we used deepTools multiBigwigSummary bins followed by deepTools plotCorrelation with arguments -- corMethod pearson, --removeOutliers, --skipZeros, --log1p.

### CTCF Differential binding analysis

To generate CTCF signal matrix used in the analysis of differential binding we used deepTools’ multiBigwigSummary BED-file, the bed file generated by merging both consensus peak sets of CTCF CUT&RUN experiments (in wild type and *Chd4*^-/-^ mice) and the drosophila normalized CTCF bigwig files (see ‘Alignment, filtering and generation of coverage track files (bigwig)’).

Differential binding was tested using DESeq2. Size factors were set to 1 for all samples (given they had already been normalized using drosophila). Default arguments were used for dispersion estimation and model building (DESeq function), and to obtain results table (results function). Results with an adjusted p-value < 0.05 were considered for further analysis.

### Over representation analysis (ORA)

Overrepresentation analysis was done using R package ‘grpofiler2’ (Kolberg L, Raudvere U, Kuzmin I, Vilo J, Peterson H (2020). “gprofiler2-an R package for gene list functional enrichment analysis and namespace conversion toolset g:Profiler.” _F1000Research_, *9 (ELIXIR)*(709). R package, v 0.2.3), using function gost with arguments ordered_query = F, correction_method = c(“gSCS”), domain_scope = “custom_annotated”, custom_bg = background_genes, using all genes detected in the single cell as background genes.

### Single cell RNA-seq

#### Preparation of single cell suspension

Testes obtained from 60 days old mice were detunicated, seminiferous tubules were gently stretched and incubated in 4 ml of DMEM culture media supplemented with 0.5 mg/ml collagenase I on an orbital shaker (60 rpm) for 10 min at room temperature. Seminiferous tubules were allowed to settle by gravity, and the supernatant containing the collagenase solution was removed. Seminiferous tubules were washed with 10 ml of DMEM by gently inverting the 15 ml tube 10 times. After the wash, the medium was discarded, and tubules were resuspended in 3 ml of fresh DMEM containing collagenase I was added together with 12.5 U of DNase I (stock concentration 1500 U/ml). Tubules were incubated on an orbital shaker for 5 min at room temperature. Subsequently, 25 μl of a 25 mg/ml trypsin solution was added and incubated for an additional 10 min. To enhance cell dissociation, an additional 8.5 μL of DNase I was added, followed by another 25 μL of trypsin, with additional incubation periods of 5 and 10 minutes, respectively. The suspension was gently pipetted five times using a trimmed pipette tip until turbidity of the media indicated release of cells. Trypsin was quenched by adding 1 ml 5% FBS and incubating for 10 min. Cell aggregates were removed by passing the cell suspension through a pre-wetted 40 μm cell strainer (mesh rinsed with DMEM). To minimize mechanical stress, portions of the suspension were filtered sequentially with gentle pipetting until the entire volume had passed through the strainer. Residual debris was reduced by dispensing the single cell suspension onto 40 ml of 0.4 % BSA in 1 × PBS to create phase separation, followed by centrifugation at 200 × g for 5 min at room temperature. The supernatant was removed, and the pellet was resuspended in 1 ml 0.04 % BSA in 1 × PBS. The suspension was then layered onto 14 ml 0.04 % BSA in 1 × PBS and centrifuged at 200 × g for 5 min at room temperature. Cells were resuspended in 1 ml 0.04 % BSA in 1 × PBS and counted in duplicate using Trypan Blue (1:1, dye:cell suspension) to confirm ≥80% viability.

The single cell suspension was further processed using Cell Multiplexing Oligo (CMO) Labeling corresponding to the single cell RNA sequencing protocol (10 x Genomics). 2 x 10^6^ cells were transferred to a 2 ml DNA LoBinding tube (cat. 022431048), and 0.04 % BSA in 1 x PBS was added for a total volume of 1 ml and centrifuged in a horizontal rotator at 200 x g for 5 min at 4 °C. The supernatant was removed, cells were resuspended in 100 μL CMO solution by gentle pipetting (10-15 times), and the suspension was incubated for 10 min at room temperature. Chilled 1 % BSA in 1 x PBS was added to the total volume of 2 ml, and the sample was centrifuged at 200 x g for 5 min at 4 °C. Cells were washed twice by removing the supernatant, resuspending the pellet in 2 ml chilled 1% BSA in 1× PBS, and centrifuging in a horizontal rotor at 200 × g for 5 min at 4 °C. Cells were finally resuspended in 200 μl chilled 0.04 % BSA in 1 x PBS and counted. Cell suspensions were adjusted with 0.04 % BSA in 1 × PBS to 1,000 cells/μl, combined at equal cell concentrations (1:1 as applicable) to a final volume of 100 μl, and kept on ice for downstream processing.

#### Alignment and data pre-processing

Alignment and initial data pre-processing was done using 10X Genomics’ Cell Ranger ‘multi’ pipeline (v 7.0.0) and ‘refdata-cellranger-arc-mm10-2020-A-2.0.0’ reference genome. Nuclear fraction of each cell was obtained using R’s package DropletQC (https://doi.org/10.1186/s13059-021-02547-0) plus the corresponding bam files generated by Cell Ranger. Further processing was done using R (v 4.5.0), tidyverse packages (https://joss.theoj.org/papers/10.21105/joss.01686) and R’s package Seurat (v 5.3.0, doi:10.1038/s41587-023-01767-y, doi:10.1016/j.cell.2021.04.048, doi:10.1016/j.cell.2019.05.031, doi:10.1038/nbt.4096, doi:10.1038/nbt.3192). Cell Ranger’s CMO-demultiplexed raw output matrixes were used to create Seurat objects.

The following per-cell metrics were also calculated in order to asses cell quality: mitoRatio, defined as the percentage of genes that are mitochondrial; logGenesPerUMI, defined as the log_10_(gene count)/log_10_(UMI count). Low quality cells were filtered out by grouping them by genotype and excluding cells that satisfied any of the following criteria: 1) mitoRatio above quantile 95, 2) nuclear fraction above quantile 95, 3) logGenesPerUMI above quantile 95, 4) logGenesPerUMI above quantile 66 and mitoRatio above 6% or below 3%, 5) logGenesPerUMI above quantile 66 and nuclear fraction above 0.3 or below 0.1. This strategy resulted in the exclusion of 34% of cells for both genotypes.

Next, for genotype-grouped cells, counts were normalized using normalizeData (scaled to the mean of counts per cell) and centered and scaled variable features (FindVariableFeatures and ScaleData) were used for PCA analysis (RunPCA (npcs = 50)). Seurat layers (corresponding to genotypes) were integrated using canonical correlation analysis (CCA) through IntegrateLayers function. Cells were clustered using FindNeighbours (using top 30 dimensions from CCA output) and FindClusters (using a resolution of 0.35).

On a first iteration, main type of cells in each cluster were broadly identified by evaluating the expression of typical marker genes. One cluster expressed no typical marker for any germ cell or somatic cell and had a low UMI count per cell and hence was discarded. All the procedure from count normalization to clustering (this time at resolution 0.7) was repeated for this new set of cells. The product of such procedure constituted the final dataset.

### Reprocessing of early germ cells subset

Germs cells from cluster GC_A (spermatogonia) to GC_C (pachytene) were selected and reprocessed: layers were re-split according to genotype, and data were reprocessed as before (from data normalization up to integration and unsupervised clustering). For visualization purposes, UMAP (RunUMAP) was run on CCA-integrated data (using top 30 dimensions and default values) to obtain a two dimension representation of the data.

### Cluster markers

To obtain cluster markers we used Seurat’s FindAllMarkers function with default parameters.

### Differential gene expression: *Chd4*^-/-^ versus wild type

To obtain lists of differentially expressed genes we used DESeq2 (v1.49.2, https://doi.org/10.1186/s13059-014-0550-8) on pseudobulk expression profiles. Pseudobulk expression was obtained using Seurat’s AggregateExpression on data grouped by mouse ID and cluster. Differential expression was obtained using the following arguments to the DESeq function from DESeq2 package: test = “LRT”, reduced = ∼ 1, fitType = “parametric”, sfType = “poscounts”, betaPrior = F.

### Processing of Chen *et al.* (2018) data set for label transfer

GSE107644_RAW.tar file was downloaded from GEO (accession number GSE107644). From all the UMI count files present in original tar file, only those annotated to contain only one type of wild type cell were used (information was extracted from file GSE107644_barcode_information.txt). The original nomenclature for some cells was changed: ‘ ln_TypeBS ‘, ‘ TypeBG2M’, and ‘G1’ were annotated as ‘spermatogonia B’; ‘ePL’, ‘mPL’, and ‘lPL’ were annotated as ‘preleptotene. A seurat object containing all original matrixes joined in a single layer were processed by normalizing data (NormalizeData), finding variable features (FindVariableFeatures), scaling data (ScaleData) and running a PCA (RunPCA).

### Label transfer: full data set as query

Since Chen *et al.* (2018) reference data set contains only up to rounded spermatids, and our dataset contains up to elongated, before conducting the label transfer protocol we first discarded cells from last 2 clusters which roughly correspond to elongating cells. We then found matching anchor cells using FindTransferAnchors (with ‘dims = 20’ and ‘reference.reduction = “pca”’). Transfer label protocol was performed with Seurat’s TransferData function using default values.

### Label transfer: spermatogonia-pachytene set as query

For our dataset consisting of spermatogonia to pachytene cells, before performing the label transfer protocol, we selected cells from spermatogonia up to late pachytene in the Chen *et al. (2018)* dataset. For this, new set of reference cells were analyzed in PCA (FindVariableFeatures, ScaleData and RunPCA(npcs = 30)). We then searched for anchor cell pairs and performed label transfer as described above.

### List of genes involved in wild type transitions between consecutive clusters

To obtain a list of genes differentially expressed between consecutive clusters in wild type cells, a similar strategy to the one described above for *Chd4*^-/-^ mice was used. All *Chd4^−/−^* cells were discarded prior to analysis. Pseudobulk expression values were obtained using AggregateExpression on data grouped by mouse ID and cluster. DESeqDataSet object was built using DESeqDataSetFromMatrix (’∼ mouse + cluster’) and differential expression results were obtained using DESeq function (test = “LRT”, reduced = ∼ mouse, fitType = “parametric”, sfType = “poscounts”, betaPrior = F).

### Statistical testing

All statistical analyses were performed in R (v 4.5.0). Associations between categorical variables were assessed using Fisher’s exact test (fisher.test, two-sided) or the chisquared test (chisq.test, with default parameters). Differences in distributions between groups were evaluated with the Wilcoxon rank-sum when two groups were compared, or Kruskal–Wallis test when more than two groups were compared (kruskal.test, default parameters), followed by post-hoc pairwise Wilcoxon rank-sum tests with Benjamini– Hochberg correction for multiple comparisons (pairwise.wilcox.test, with arguments p.adjust.method = “BH”).

When applicable, multiple hypothesis testing was controlled using the Benjamini– Hochberg procedure.

## Acknowledgments

This study was supported by NIH/NICHD (grant HD110990), and PHF (grant 4431-10071).

