## Supplementary figure legends for "A novel CHD4-CTCF regulatory axis drives key developmental programs and transitions in male germ cells"

**Supplementary Figures**

**Figure S1. Characterization of Chd4^−/−^** **mice. (A)** TUNEL-DAPI stained histological sections (left) and it's quantification (right) of wild type (1.87 ± 1.06 apoptotic germ cells per tubule with at least one apoptotic cell, n = 62 seminiferous tubules) and *Chd4^−/−^* (4.68 ± 3.21, n = 142 seminiferous tubules; P < 0.0001, two-tailed t test) testis. **(B)** Composition of prophase I spermatocyte population in *Chd4^−/−^* and wild type mice. **(C)** H&E-stained histological sections of wild type and *Chd4^−/−^* ovaries. F, follicles and CL, corpora lutea. **(D)** Representative images of wild type and Chd4^−/−^ spermatocytes immunostained with SYCP3 and SYCP1 or SYCE1.

### **Figure S2. Cluster and genotype-wise quality control metrics for cells in scRNA-seq full set.** All metrics are computed per cell. Nuclear fraction denotes the proportion of RNA molecules containing introns. mitoRatio represents the percentage of reads mapping to mitochondrial genes. Genes refers to the number of detected genes, and UMIs to the number of unique RNA molecules detected.

### **Figure S3. ScRNA-seq of wild type and *Chd4^−/−^* testes. (A)** Clustering of full scRNA-seq experiment. The full set of cells analyzed in the scRNA-seq experiment shown in UMAP space with their respective clustering, grouped by genotype. **(B)** Cell label transfer using the dataset from Chen *et al*., 2018. The cells shown in panel A were assigned an identity based on label transfer using the dataset from Chen et al, 2018 as reference. This dataset used as reference contains from spermatogonia A1 up to rounded cells, hence, cell stages past round spermatids in our dataset have no label. Cells' labels are as follows: spermatogonia A1 ('A1'), Intermediate spermatogonia ('In'), spermatogonia B ('B'), preleptotene ('PL'), leptotene ('L'), zygotene ('Z'), early pachytene ('eP'), middle pachytene ('mP'), late pachytene ('lP'), first meiotic division (MI), second meiotic division (MII), round spermatids (steps 1 and 2 'RS1o2', steps 3 and 4 'RS3o4', steps 5 and 6 'RS5o6', steps 7 and 8 'RS7o8'). **(C)** Number of cells per cluster and genotype (full dataset). Plot is divided in two: on the left are germ cells, on the right are somatic cells. **(D)** Heatmap of label transfer results using the Chen et al, 2018 as reference (x-axis) and unsupervised clustering of our full dataset (y-axis). Each cell of the matrix shows the number of cells assigned to a given reference label within each unsupervised cluster. Color intensity represents the number of cells. Clusters with fewer than 10 cells are not annotated with numbers. **(E)** Expression of marker genes per cluster for the full scRNA-seq dataset. Dot plot depicting the scaled average expression (z-score) of selected marker genes across clusters. Dot size represents the percentage of cells expressing each gene within the cluster. **(F)** Heatmap of label transfer results using the Chen et al, 2018 as reference (x-axis) and unsupervised clustering of our spermatogonia-pachytene dataset (y-axis). Each cell of the matrix shows the number of cells assigned to a given reference label within each unsupervised cluster. Color intensity represents the number of cells. Clusters with fewer than 10 cells are not annotated with numbers.

**Figure S4. CHD4 genome binding experiments. (A-B)** Left: signal correlation across the whole genome divided in bins of 10 kilo bases for spermatogonia **(A)** or preleptotene **(B)** genome binding experiments. Pearson correlation coefficient is indicated inside each cell. Right: Euler diagram showing intersection of peaks for the mentioned experiments. **(C)** Intersection of consensus peaks for spermatogonia and preleptotene CHD4 genome binding experiments.

**Figure S5. CHD4 is associated with genes that change upon differentiation.** Left **(A-B):** Euler diagram showing overlap between genes that are differentially expressed in wild type cells when comparing cluster GC_1 and GC_2 (A) or GC_2 and GC_3 (B) ('Genes that change in the transition'), genes annotated with a CHD4 peak that are differentially expressed when comparing wild type and *Chd4^−/−^* cells in either GC_1 or GC_2 (A) or in GC_2 or GC_3 (B) ('Genes with peak DE in either cluster'), among all genes detected in the scRNA-seq assay ('All genes'). Below Euler plot are annotated p-value and odds ratio value from Fisher's exact test, on categories 'Gene is annotated with a CHD4 peak and is differentially expressed in either cluster from transition' and 'Genes is differentially expressed in wild type cells during the transition'. Right **(A-B)**: percentage of genes annotated with a CHD4 peak that are differentially expressed in *Chd4^−/−^* cells (compared to wild type) in either of the clusters from the transition (GC_1 or GC_2 (A), GC_2 or GC_3 (B)), among genes that either are differentially expressed in wild type cells during the transition ('GC_1→GC_2' (A), 'GC_2→GC_3' (B))) or not ('Not GC_1→GC_2' (A), 'Not GC_2→GC_3' (B)).

### **Figure S6. CHD4 modulates the transcription of genes involved both in spermatogonia differentiation and in transition to meiosis. (A–C)** Normalized mean expression profiles of individual genes across clusters GC_1–GC_5 in wild type (WT) and *Chd4^−/−^* cells. Each line corresponds to one gene. For visualization, combined WT+ *Chd4^−/−^* profiles were clustered using k-means (k = 2), and results are shown as “Subset 1” and “Subset 2”. **(A)** Genes from the upper-left quadrant of Figure 4B (GC_1→GC_2 transition, GC_2 DEGs). **(B)** Genes from the upper-left quadrant of Figure 4D (GC_2→GC_3 transition, GC_3 DEGs). **(C)** Genes from the upper-right quadrant of Figure 4D (GC_2→GC_3 transition, GC_3 DEGs). Genes of interest are highlighted in color.

**Figure S7. CTCF genome binding experiments. (A)** Signal correlation across the whole genome divided in bins of 10 kilo bases for CTCF CUT & RUN experiments. Pearson correlation coefficient is indicated inside each cell. **(B,C)** Euler diagrams showing intersection of peaks for CTCF CUT & RUN replicates in wild type **(B)** or *Chd4^−/−^* **(C)** cells. **(D)** Intersection of consensus peaks for CTCF experiments in both genotypes.

**Figure S8. Aggregate profiles of histone marks, CHD4 and CTCF in CHD4 peaks, grouped by their distance to the nearest TSS and overlapping with CTCF.** Aggregate profiles depicting mean signal for different histone marks, CHD4 (both replicates) and CTCF (in wild type mice, both replicates), in preleptotene cells. In the upper panel, profiles are centered at CHD4 preleptotene consensus peaks that are located further away than -2000 or +1000 bp from the closest TSS (distal peaks). In the lower panel, profiles are centered at the closest TSS of those CHD4 preleptotene consensus peaks that are located within -1000 to +300 bp of a TSS. In both cases profiles are grouped by whether they overlap (green) or not (blue) a CTCF preleptotene consensus peak in wild type mice.

**Figure S9. Aggregate profiles of histone marks, CHD4 and CTCF in CTCF peaks, grouped by their distance to the nearest TSS and overlapping with CHD4**. Aggregate profiles depicting mean signal for different histone marks, CHD4 (both replicates), and CTCF (in wild type mice, both replicates), in preleptotene cells. In the upper panel, profiles are centered at CTCF preleptotene consensus peaks that are located further away than -2000 or +1000 bp from the closest TSS (distal peaks). In the lower panel, profiles are centered at the closest TSS of those CTCF preleptotene consensus peaks (in wild type mice) that are located within -1000 to +300 bp of a TSS. In both cases profiles are grouped by whether they overlap (green) or not (blue) a CHD4 preleptotene consensus peak.

**Figure S10. Aggregate profiles of histone marks, CTCF and CHD4 in CTCF peaks, grouped by their distance to the nearest TSS and their DBR status.** Aggregate profiles depicting mean signal for different histone marks, CHD4 (both replicates) and CTCF (in *Chd4^−/−^* and wild type mice, both replicates), in preleptotene cells. In the upper panel, profiles are centered at CTCF preleptotene consensus peaks that are located further away than -2000 or +1000 bp from the closest TSS (distal peaks). In the lower panel, profiles are centered at the closest TSS of those CTCF preleptotene consensus peaks (in wild type mice) that are located within -1000 to +300 bp of a TSS. In both cases profiles are grouped by their DBR status: positive DBRs (yellow), negative DBRs (blue) and non-DBR (light blue).
