## Supplementary Figure 1 for "A novel CHD4-CTCF regulatory axis drives key developmental programs and transitions in male germ cells"

### Slide 1
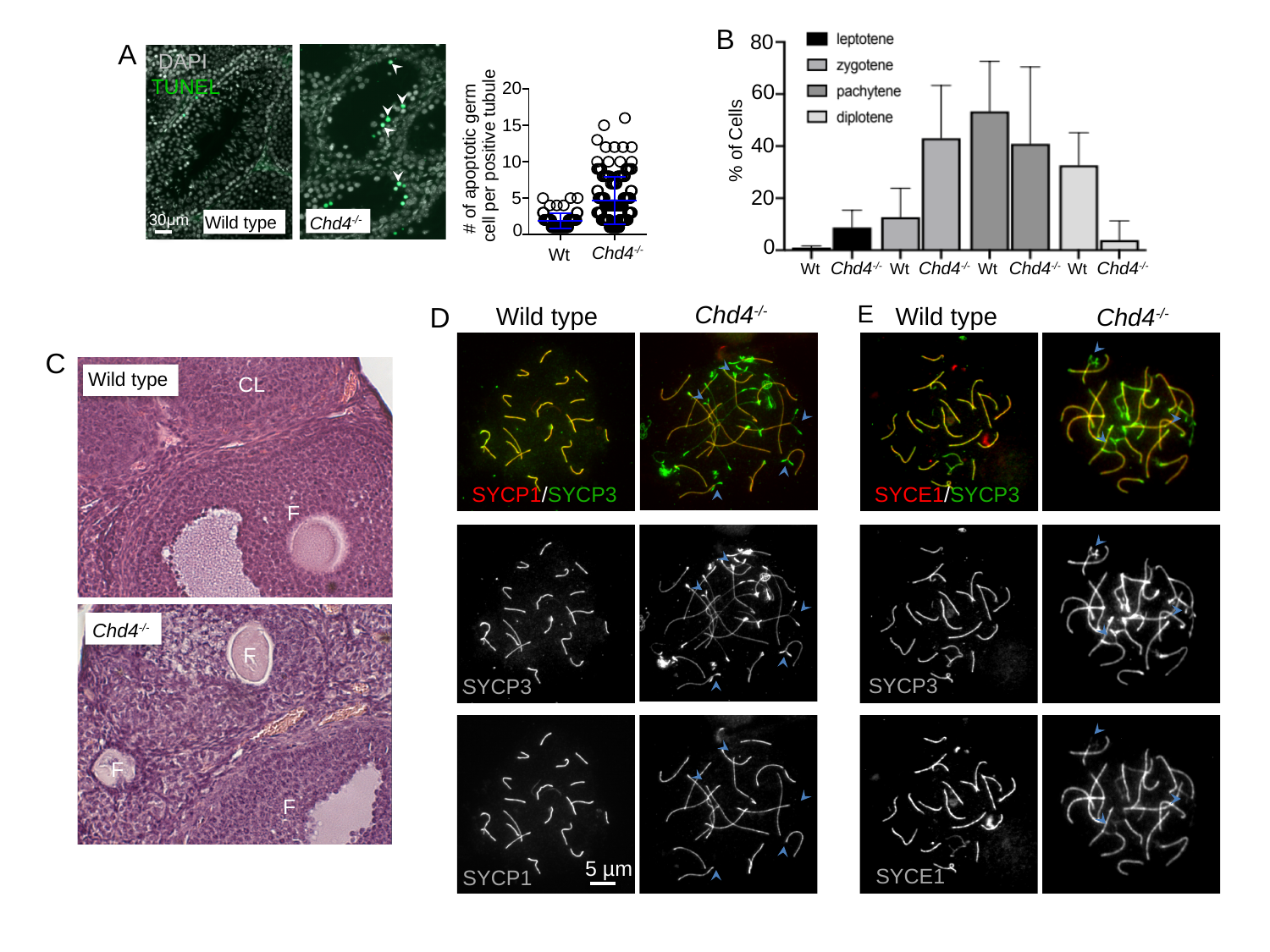

B
80
60
% of Cells
40
20
0
Wt
Chd4^-/-^
Wt
Chd4^-/-^
Wt
Chd4^-/-^
Wt
Chd4^-/-^
A
A
DAPI TUNEL
#### of apoptotic germ
cell per positive tubule
Wt
20
15
10
5
0
Chd4^-/-^
Wild type
Chd4^-/-^
30μm
E
Chd4^-/-^
D
Wild type
Wild type
Chd4^-/-^
C
Wild type
CL
SYCE1/SYCP3
SYCP1/SYCP3
F
Chd4^-/-^
F
SYCP3
SYCP3
F
F
5 µm
SYCE1
SYCP1
