## Supplementary figures and images for "A novel CHD4-CTCF regulatory axis drives key developmental programs and transitions in male germ cells"

### Supplementary Figure 2

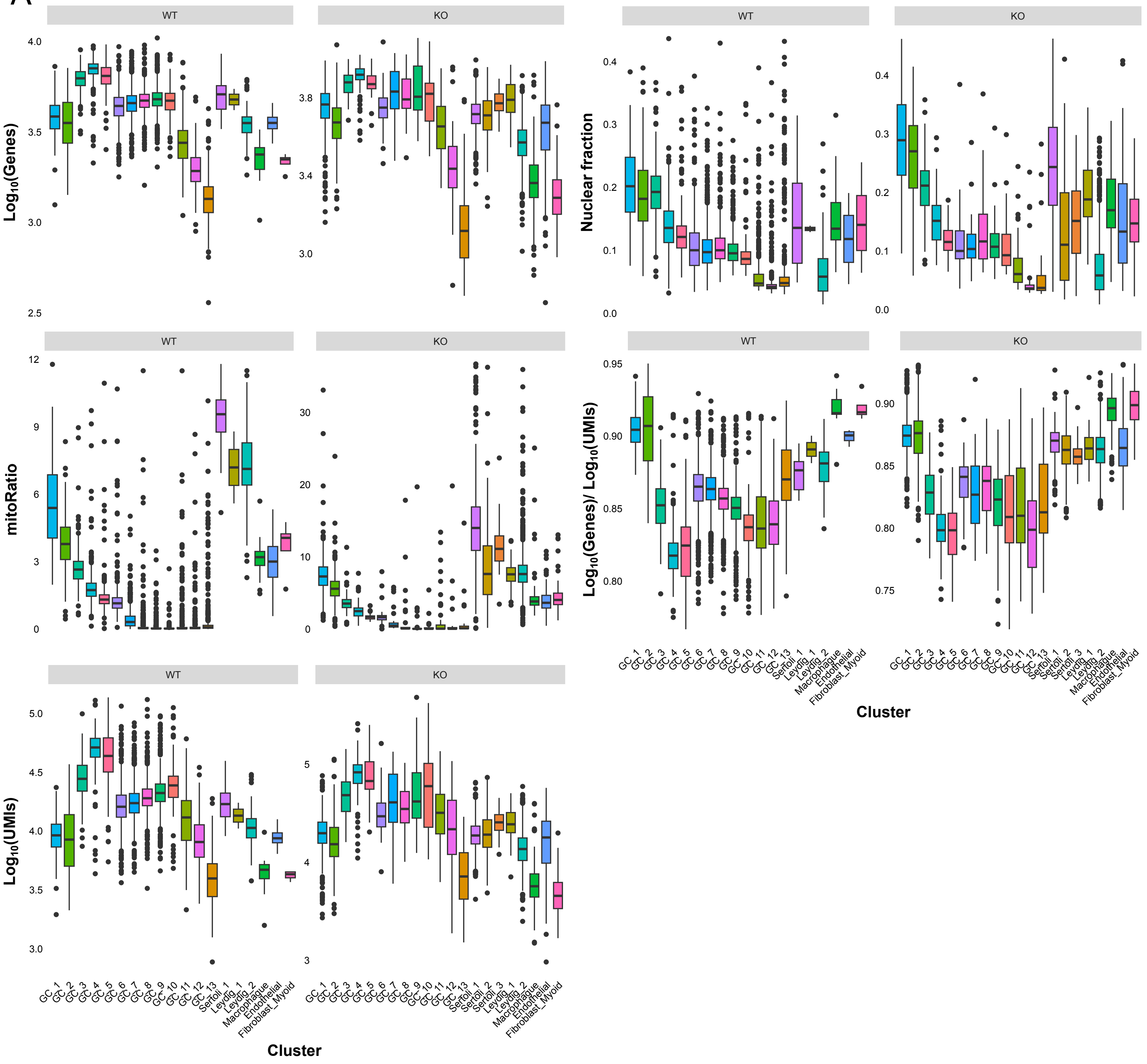

### Supplementary Figure 3

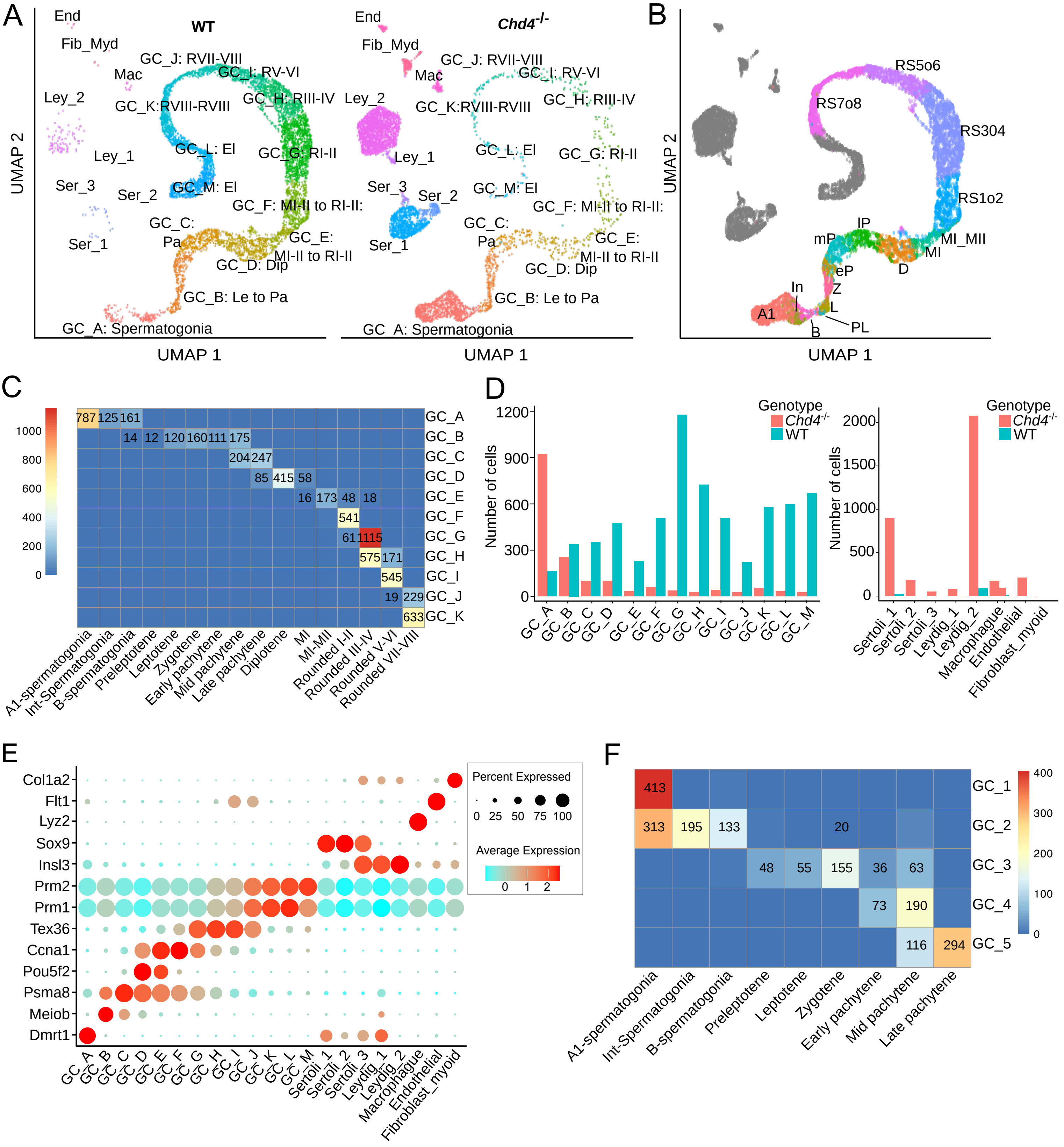

### Supplementary Figure 4

A

#### Spermatogonia

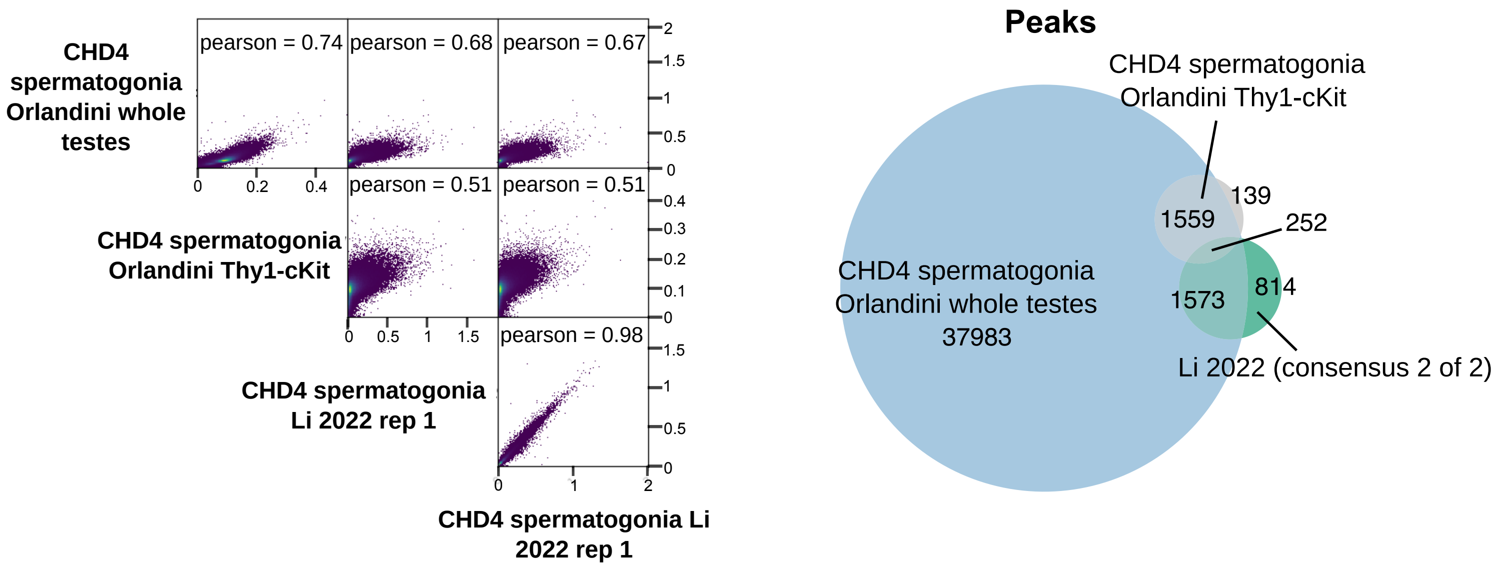


B

#### Preleptotene/leptotene

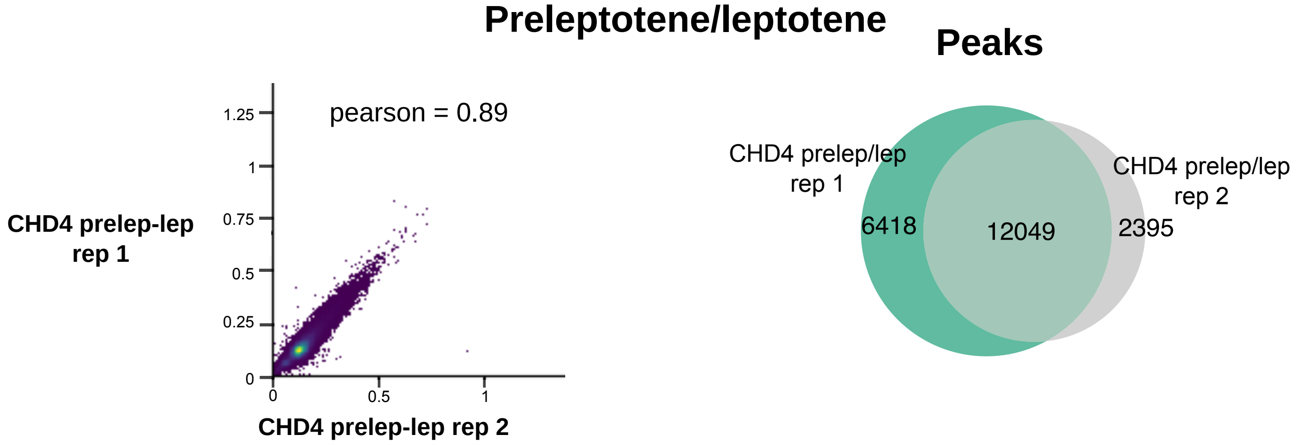


C

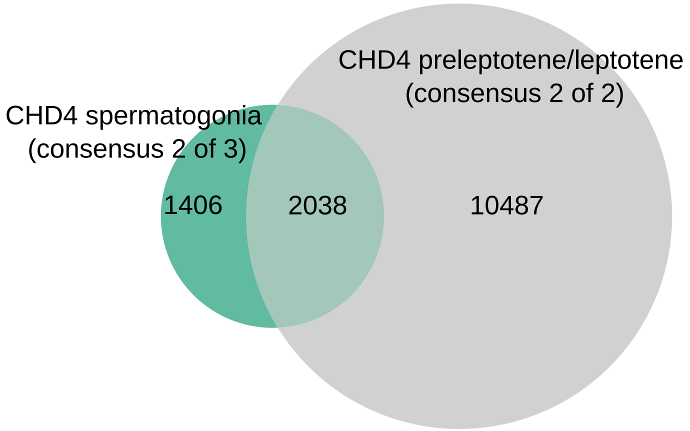

### Supplementary Figure 5

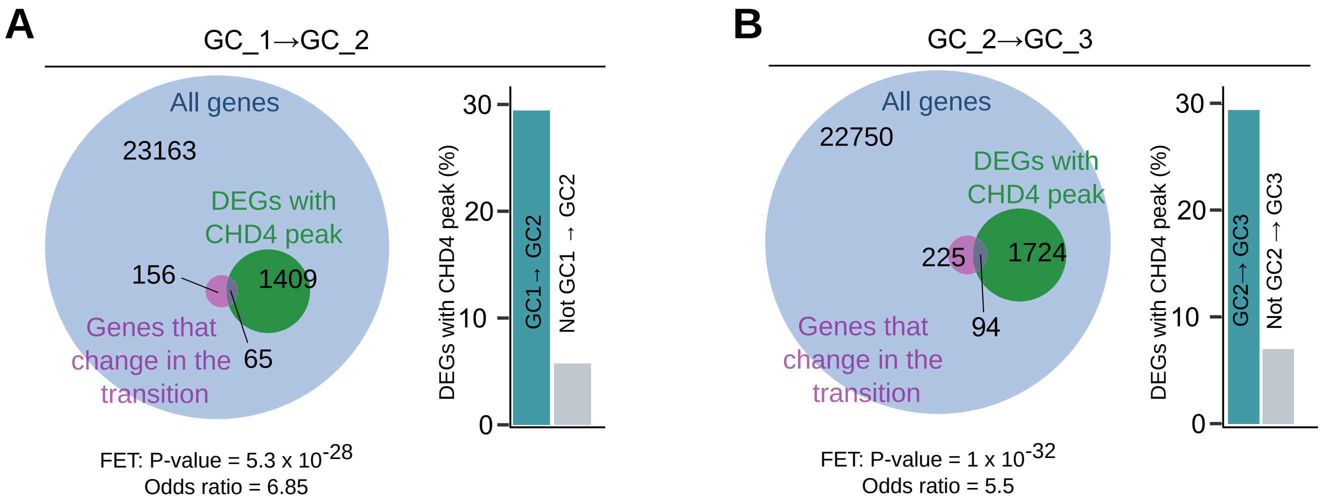

### Supplementary Figure 6

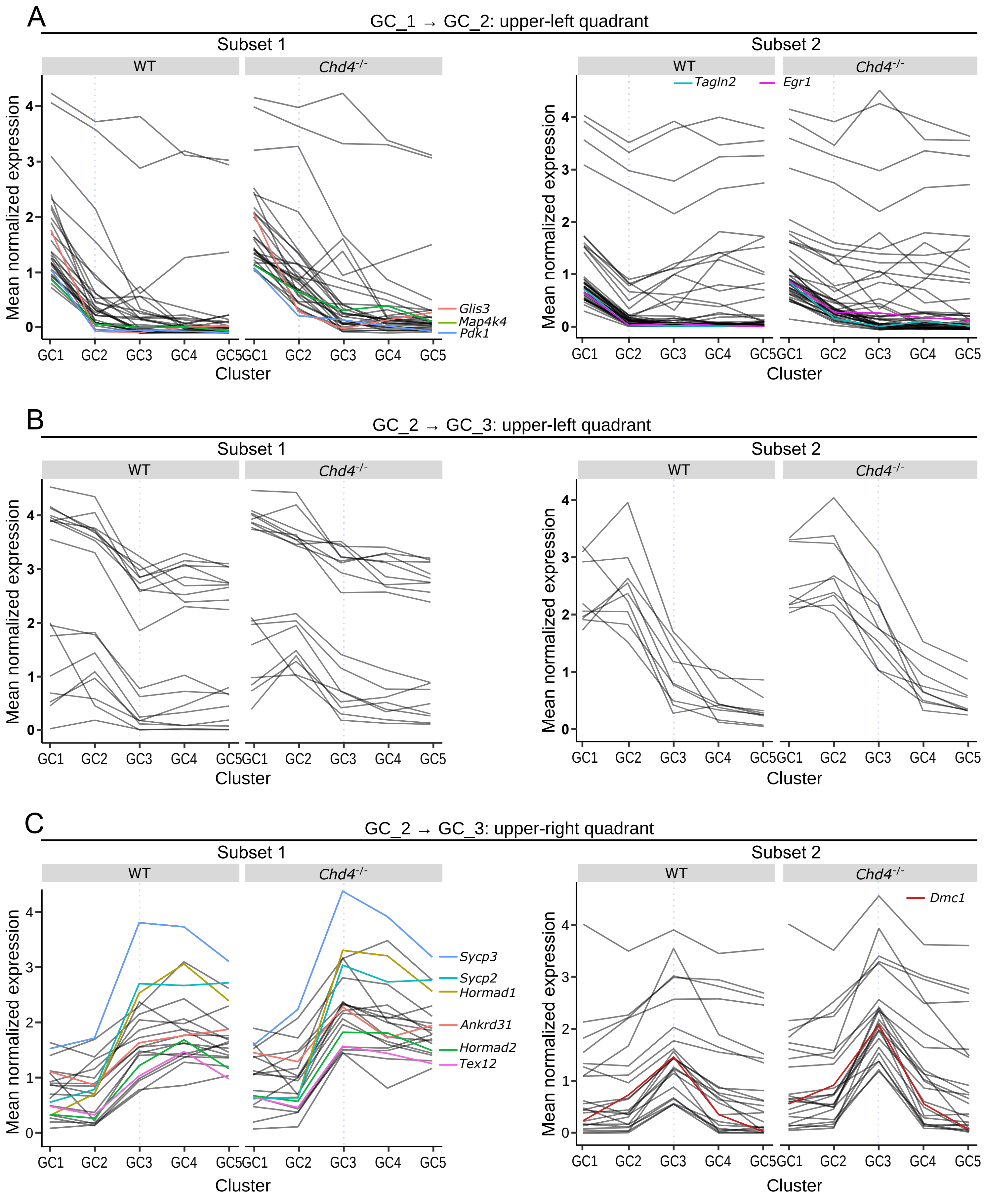

### Supplementary Figure 7

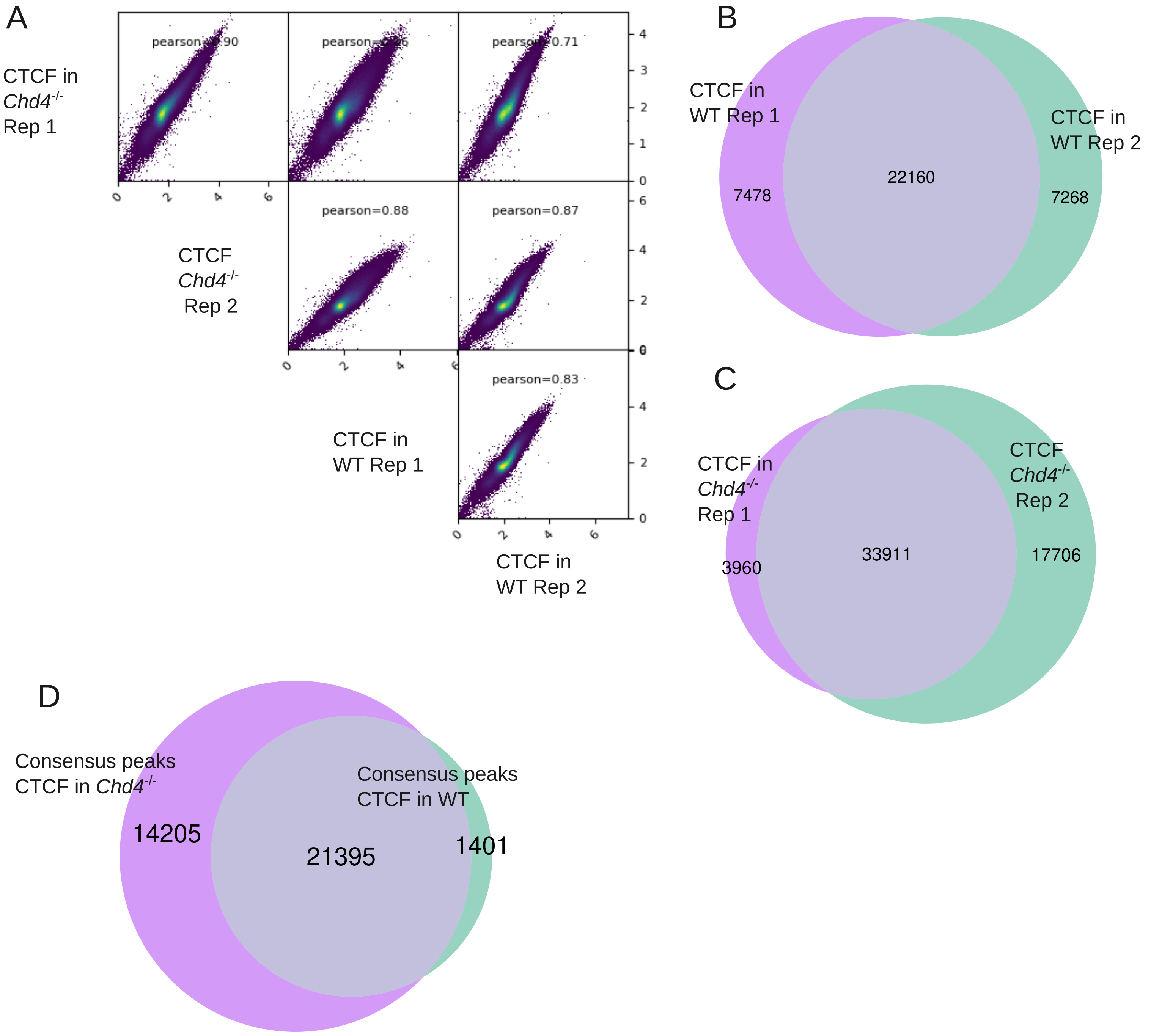

### Supplementary Figure 8

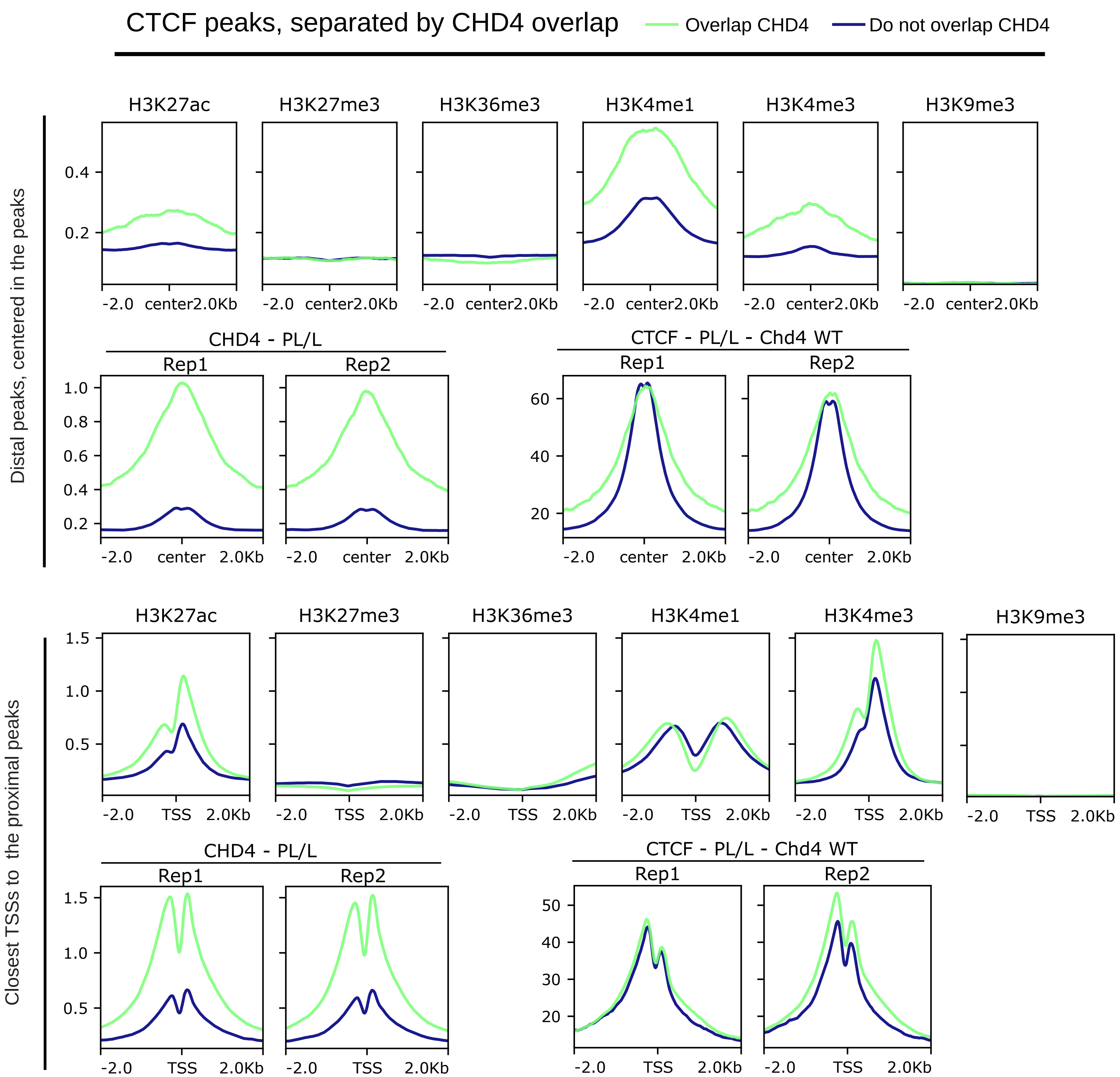

### Supplementary Figure 9

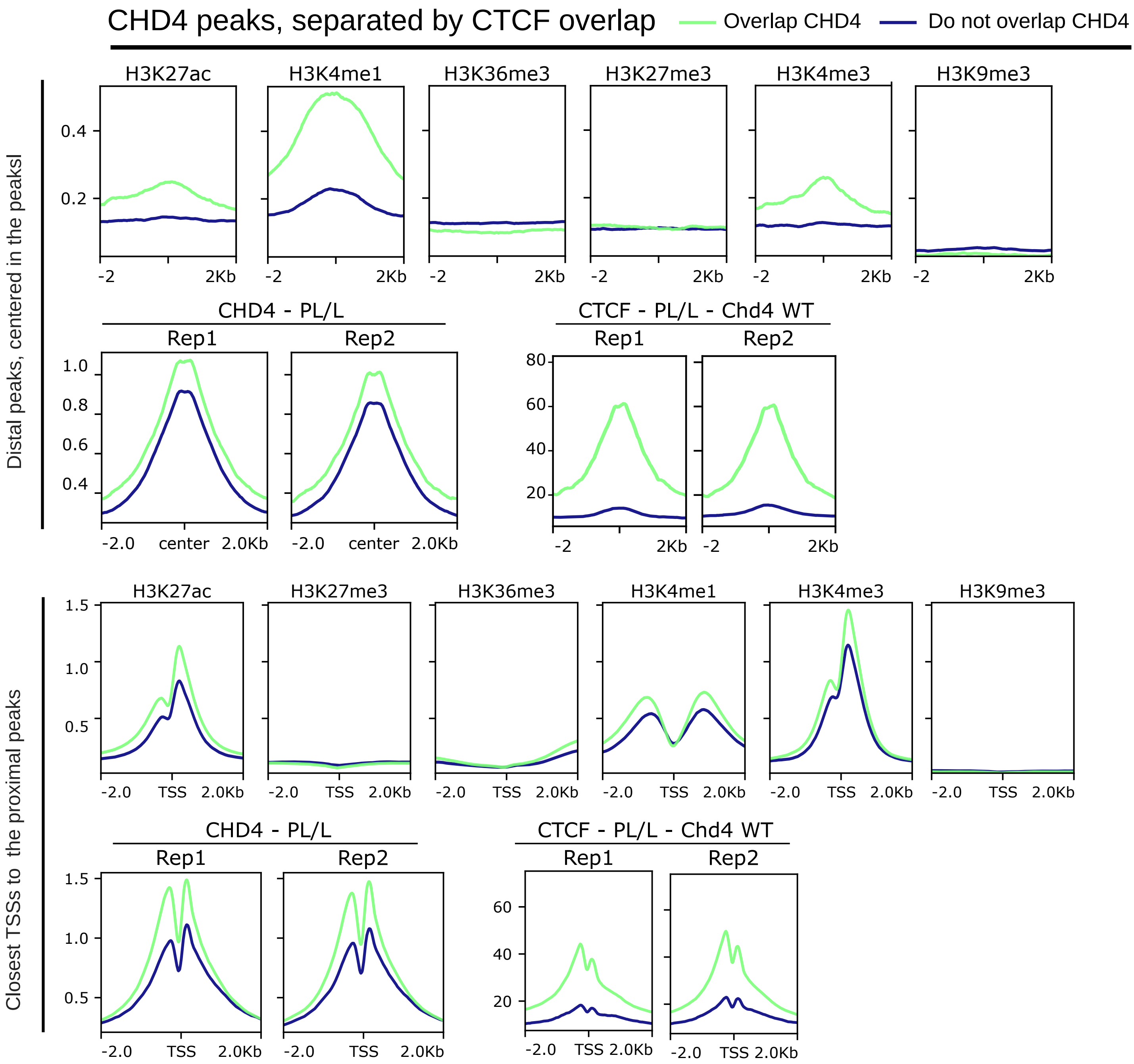

### Supplementary Figure 10

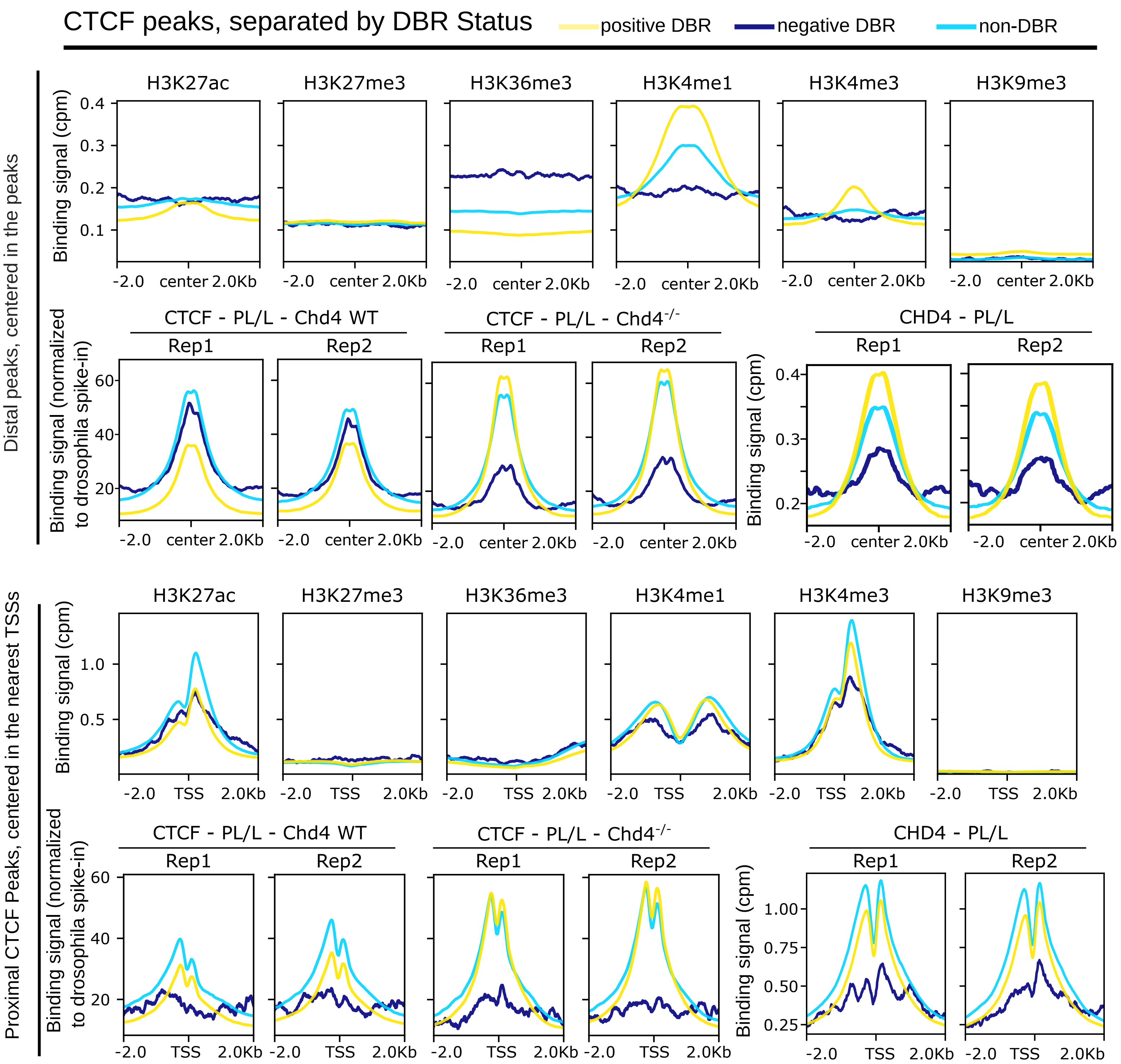
